# Quantum analysis of P3a and P3b from auditory single trial ERPs differentiates borderline personality disorder from schizophrenia

**DOI:** 10.1101/2022.09.23.509191

**Authors:** Dmitriy Melkonian, Anthony Korner, Russell Meares, Anthony Harris

## Abstract

Traditional approaches to EEG modelling use the methods of classical physics to reconstruct scalp potentials **i**n terms of explicit physical models of cortical neuron ensembles. The principal difficulty is that the multiplicity of cellular processes with an intricate array of deterministic and random factors prevents creation of consistent biophysical parameter sets. An original, empirically-testable solution has been recently achieved in our previous studies by a radical departure from the deterministic equations of classical physics to the probabilistic reasoning of quantum mechanics. This crucial step relocates elementary bioelectric sources of EEG signals from the cellular to the molecular level where positively and negatively ions are considered as elementary sources of electricity. The rationale is that despite dramatic differences in cellular machineries, statistical factors governed by the rules of central limit theorem produce EEG waveforms as a statistical aggregate of the synchronized activity of multiple closely-located microscale sources. Using the formalism of nonhomogeneous birth-and-death processes (BDP) the quantum models of microscale events are deduced and linked to the dynamics of macroscale EEG waveforms. This study expands these methods with new features for comprehensive analysis of event related potentials directly from single trials, i.e. the EEG segments which are closely related in timing to cognitive events. We derive a universal model of the components of single trial ERPs both in frequency and time domains. This, for the first time, enables us to quantify all significant cognitive components in single trial ERPs, providing an alternative to the traditional method of averaging. Given P300 as an important objective marker of psychiatric disorders, a methodology which reliably discloses the component compositions of this potential, may have specific diagnostic importance. In this study, reliable identification of the P3a and P3b components from an auditory oddball paradigm provided a means of differentiating borderline personality disorder from schizophrenia.

## Introduction

The recording of human electroencephalogram (EEG) by means of electrodes on the scalp is one of the most widely-used functional tests of neural function with excellent temporal resolution. The exquisite sensitivity of EEG to changes in mental activity has been documented in numerous studies. A fundamental criterion used to link electrophysiological and cognitive variables is a concomitant variation of neurophysiological and psychological processes. In consequence, the key to understanding of the information processing context of EEG signals is provided by detection of the changes in ongoing EEG activity which is time-locked (or at least closely related in timing) to a particular cognitive event.

The EEG peaking waveform which frequently appears at specific time intervals linked to the application of a cognitive stimulus is known as an event related potential (ERP). An overlap of ERP components with the ongoing oscillations of the EEG is a factor which obscures the morphology of ERPs. Therefore, signal extraction methods are absolutely essential in ERP analyses.

The most common method is ensemble averaging which consists in summation of a series of EEG epochs (trials), each of which is time-locked to the event of interest. The basic assumption of the method is that pertinent EEG fragments are the summands of two sources: (i) the ERPs which are constant over trials, (ii) random constituents that are not time-locked to the event. The first assumption asserts that identical time-locked ERPs are expected in response to repeated cognitive events. However, there is a great deal of experimental evidence that this assumption is an oversimplification which disregards the fact that ERP composition consists of multiple components, each of which is subject to trial-to-trial variability governed by different probabilistic laws.

The fact that the same event can elicit somewhat different signals has been evident for a few decades and is clearly seen from the following general psychophysiological definition of endogenous ERP components: “The components must be nonobligatory responses to stimuli. The same physical stimulus, presented to the same subjects, sometimes will and sometimes will not elicit the component” [1].

Consequently, the conventional ensemble average would not necessarily correspond to any of the individual single-trial responses. This means that without an account of stochastic factors, ensemble averaging creates ambiguity with respect to the analysis and interpretation of ERPs. An account of the diversity of single trial ERPs provides more information related to changes of cognitive state in response to the stimulus.

A lot of effort has been invested in inventing methods of estimating ERPs directly from single trials. The difficulty in developing this approach relates to the basic mechanisms that underlie the generation of the EEG signal.

It is generally accepted that EEG signals are distant manifestations of synchronized activities in populations of cortical neurons. The processes involved are complex and their interpretation rests mainly on an empirical understanding. The development of theoretical foundations has been widely researched and, until recently, has been approached using the methods of classical physics [2, 3]. The major proposition is that cortical neurons are the elementary microscale sources of the EEG waveforms. Such an approach assumes that, in principle, EEG dynamics can be deduced from physical models of neuronal ensembles [2]. On these grounds, the membrane potentials produced in some way by the cortical neurons appear as the “building blocks”, from which the EEG waveforms are composed. In particular, the proposition that EEG waves are constituted primarily by the postsynaptic potentials of cortical neurons is widely accepted.

However, because electrical activity associated with any particular neuron is small, it is only possible, using scalp electrodes, to detect the integrated activity of a large number of neurons. The creation of a corresponding model using the methods of the classical theory of electromagnetism would need to be supported by the parameters of all participating neurons. Significant difficulties are created by the anatomical complications posed by the multiplicity of cellular elements, along with an insufficient knowledge of their morphological details and functional relationships. This leads to an intractably large number of degrees of freedom and prevents a unique determination of mass effect. Under conditions of such uncertainty little confidence can be placed in particular solutions from a large number of different models. It is clear that no single model has yet achieved the goal of integrating the wide variety of parameters of separate neurons or neuronal ensembles into the dynamics of EEG waveforms. In a review of the state of art, Michael Cohen notes that “surprisingly little is known about how the activity in neural circuits produces the various EEG features” [4].

A radically different approach to the treatment of mass potentials, the category of signals to which the EEG and single trial ERP belong, has been achieved by a departure from the deterministic equations of classical physics to the probabilistic formalism of quantum mechanics [5]. The crucial step is relocation of the microscale origins of the macroscale potentials from a cellular to a molecular level. Instead of the continuous time membrane potentials implemented in previous theories, the key role for elementary cortical sources of electricity is attributed to ions, positively and negatively charged particles, the size and stochasticity of which conform to quantum mechanics.

A general solution is supported by a particle model using non-homogenous birth and death processes (BDP) for description of trans-membrane transport of ions. A crucial outcome is a link between the global scale mass potential and the underlying microscale events. This link provides a way of defining the universal objects of mass potential, both on the micro- and macro-scales. On the global scale a universal object called the half-wave function (HWF) is associated with positive- and negative-peaking waveforms of the mass potential. The microscopic scale counterpart of the HWF is described as a transient amalgamation of deterministic and stochastic processes called the transient deterministic chaos. Thus, quantum theory puts models of various types of mass potentials on a common theoretical and computational basis.

The quantum theory of mass potentials is subsequently elaborated in a number of ways for EEG and single trial ERP analyses [6]. Computational algorithms in these applications include the method of high-resolution fragmentary decomposition [7] which decomposes EEG segments, including single trial ERPs, into the sums of HWFs.

The processing of characteristic EEG and single trial ERP recordings has shown that this novel methodology effectively resolves component overlap. An appealing feature of this innovation is the creation of adequate models of complex ERPs, made up of several overlapping sub-components. The remarkable accuracy of such procedures is demonstrated using typical single trial ERPs from healthy subjects.

The purpose of this study is to apply the quantum theory introduced in previous papers [5,6], to the analysis of single trial ERPs. While the first part deals mainly with fundamentals, it is followed by an application to psychiatric disorders, in the form of a comprehensive comparative analysis of the auditory P3a and P3b from single trial ERPs in patients with borderline personality disorder (BPD) and schizophrenia. Both illnesses are disorders of integration [8]. Though the form of disintegration in the two conditions differs functionally, the pattern of symptoms can sometimes overlap. For example, both may experience hallucinations [9]. Misdiagnosed patients may risk being exposed to the wrong treatment.

The attempts to improve diagnostics using electrophysiological tools have been focused on the P3 event-related potential (ERP). The P3 has been broadly studied in psychiatric disorders and is widely accepted as the most important marker of cognitive functions [1]. Reduction in amplitude of the average P3 from the standard auditory oddball paradigm is one of the most replicable biological observations of schizophrenia, present regardless of medication status [10]. A similar abnormality in BPD has been reported in a comparative study of patients with BPD and schizophrenia, where the authors report, “The ERP abnormalities found in patients with BPD are indistinguishable from those found in patients with schizophrenia” [11]. However, the methodology of average ERPs used in the study ignores the complex component composition of P3, specifically its P3a and P3b sub-components, reflecting contributions from various generators.

The importance of the account of component composition of P3 has been demonstrated in the study of single trial auditory ERPs in BPD patients [12]. The P3a, which depends on the circuitry having prefrontal connections, was significantly larger than in normal controls and similar in amplitude to young adolescents. In addition, the P3a failed to habituate, which suggested a failure of the pre-frontally mediated inhibitory mechanisms. It is widely acknowledged that prefrontal deficiencies are also evident in schizophrenia.

These facts indicate a need, unfulfilled by traditional methodologies of average ERPs, to enhance the comparative analysis of BPD and schizophrenia by reliable identification of the dynamics of single trial P3a and P3b as well as the trial-to-trial variability of these components.

## Theory

### Understanding EEGs at the molecular level: the “Stion”

It is generally assumed that EEGs are distant manifestations of electrical phenomena occurring on the microscopic scale, consisting of cortical ensembles of multiple excitable cells immersed in interstitial fluid. This activity can be recorded on the surface of the scalp because the tissue that lies between the source and the scalp acts as a volume-conductor.

The elementary sources of the EEG are ions, both positively and negatively charged particles, which cross the cell membranes in both directions. The probabilistic nature of these microscopic scale events was discovered by the patch-clamp technique, which provides a means of measuring ion currents through individual ion channels in a cellular membrane [13]. The fundamental finding was that individual ion channels are essentially stochastic entities that open and close in a random way [13]. Thus, the behavior of ions is governed by probabilistic rules.

The contribution of a single ion to the changes in electrical potential differences between various locations in the extracellular space is vanishingly small. This means that measurable changes of the extracellular potentials (field potentials) are produced by the mass effects of multiple elementary sources [14]. Two requirements must be fulfilled for this integration to occur: a) the cells must be closely located and comprise an ensemble in order to have functional connectivity; (b) the activation of the cells must be synchronized.

The interior and exterior of a cell, be it a neuron or a glial cell, are both varieties of saline solution (water with ions dissolved in it) separated by membranes. Considering the membrane as a border, tissue can be divided into an extracellular and intracellular space. To a good approximation, the extracellular space can be considered independent of the intracellular space, because its boundaries, the cell membranes, have high resistances (several kΩ·cm^2^), compared with the resistance of the extracellular space (~200 Ω·cm). An important factor is that, within the range of frequencies of physiological interest (0-1kHz), the capacitive, inductive, magnetic, and propagative effects of bioelectrical phenomena in the extracellular space can be neglected [15]. Thus, the extracellular space may be regarded with reasonable accuracy as a resistive medium.

The populations of ions separated by membranes are illustrated by the schematic diagram in Fig 1. The whole-colored cloud in the left panel shows a cell ensemble which is capable of producing changes in the field potentials. We call it a local cortical generator (LCG). The ions inside the cells of the LCG are considered as contents of the interior compartment **I,** shown by the circle. The ions from the exterior of the cells composing the LCG are considered as the contents of the external compartment **E**.

**Fig 1.**
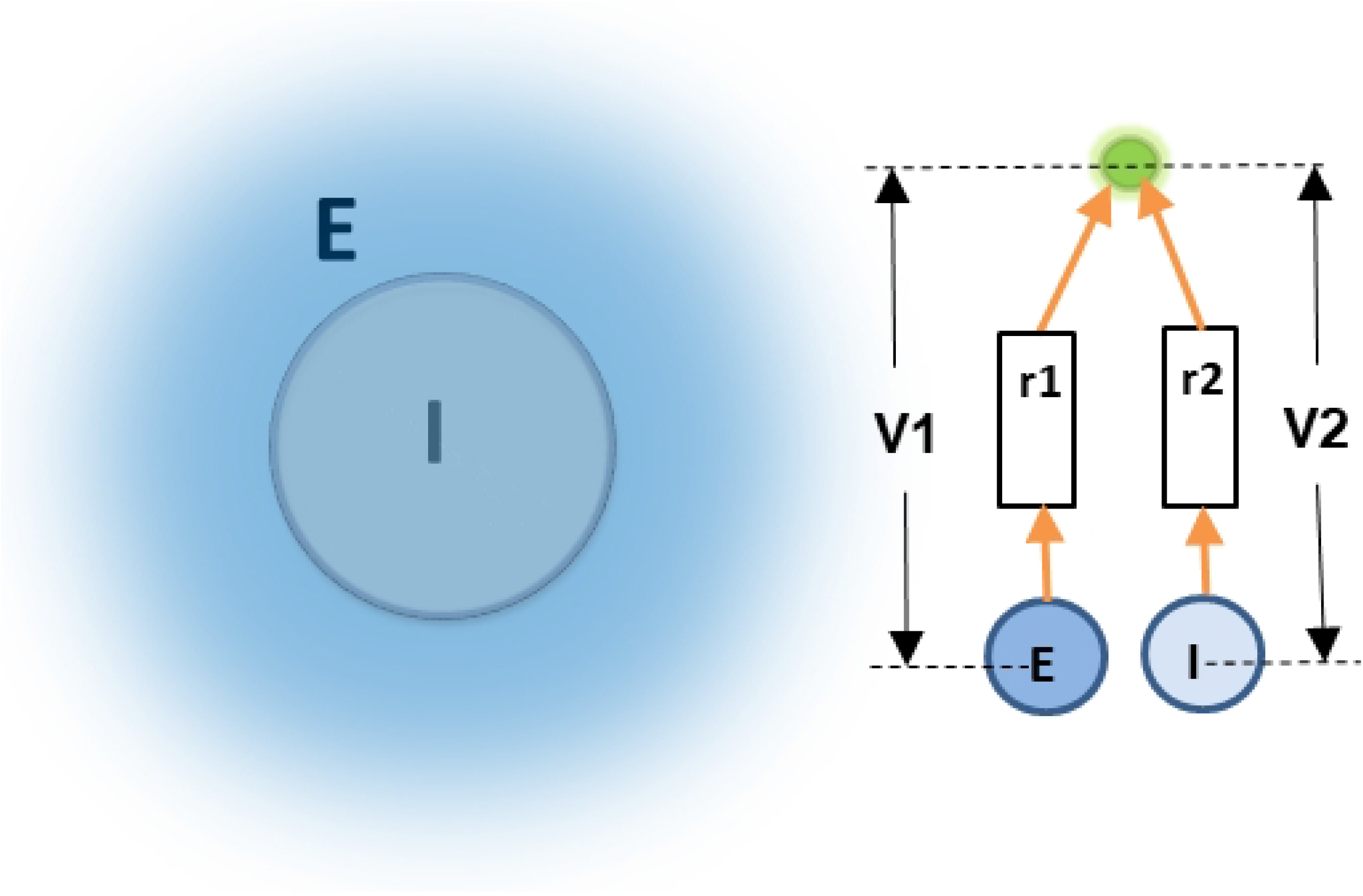
Schematic diagram of LCG and its E and I compartments.

Since the **I** and **E** compartments are separated by membranes working as insulators, they are shown in the right panel of Fig 1 as isolated, separate sources of electricity, **E** and **I**, respectively.

To evaluate the contribution of ions from the E and I compartments to the generation of field potentials, we consider the potential differences V1 and V2 between the compartments and electrical ground shown in the top of the right panel as a small green circle. This element may be associated with the cortical electrode.

In fact, it is not the voltages themselves but electric currents flowing in the extracellular space that define the influence of the LCG on field potentials. To a good approximation these currents are defined as i1=V1/r1 and i2=V2/r2. Here r1 is the resistance of the extracellular space, while r2=r1+rM, where rM is the resistance of the membrane. The internal structure of such systems may be almost impossible to delineate precisely. However, we do not need to deal with detailed descriptions of specific circuits. The general framework of our theory, which is not particularly concerned with the morphology of the LCG, is the fact that rM » *r*1. Accordingly, i1 » *i*2. Separating the interior of the cells from the extracellular space, membranes prevent the ions located inside the neurons from producing measurable changes of the current flow in the extracellular space. In contrast, the cumulative effects of the charges of the ions released from the cells during synchronized activation of cellular ensembles do influence the dynamics of the global scale EEG.

On these grounds the global scale potential is linked to the net charge of positive and negative ions released from the cells of the LCG. We have termed such an ensemble of ions a STION (STochastic IONs), in an earlier paper [6].

Physically, the population of particles considered as a STION represents a thin cloud of cations and anions spread over the outer surfaces of the membranes of the cells composing the LCG (closely located and functionally-linked neurons). During resting conditions, the transmembrane ion transport is balanced. This means that, given a local volume, the numbers of positive and negative particles randomly fluctuate over the mean values. Under these conditions the STION does not produce measurable changes of the macroscale voltages. The situation is changed under the influence of the triggering event which induces transient exchange of the ions between the **I** and **E** compartments. The transport of ions changes the contents of both the E and I compartments. Due to the high electrical resistance of membranes, the changes of the **I** compartment do not produce detectable changes of the current flow in the extracellular medium. In contrast, the **E** compartment behaves as a STION: an ensemble of extracellular particles which define the flow of currents in the extracellular medium.

### Half-wave function

There are several probabilistic methods within quantum theory, an essential goal of which is to describe the behavior of multi-particle systems with many degrees of freedom in terms of global systems with only a few “macroscopic” degrees of freedom [16]. The realization of such approaches needs to be supported by an adequate probabilistic model. In this context the reference is made to the central limit theorem as a rule that defines the limiting behavior of ensembles of random variables [17]. This theorem states that the sum of a large number of random samples from independent sources always tends to the limit in the form of a normal distribution.

Let n charged particles comprising the STION be described by the random variables g_1_, g_2_,…,g_n_ with means η_1_,η_2_,…,η_n_, and variances, 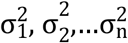. With an increasing number of particles (i.e. 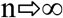), the limiting behavior of their exponential Fourier transforms G_1_(iω),G_2_ (iω),…,G_n_(iω),… converges under quite general conditions to the limit probability distribution in the form of the following function of complex variables [18]:

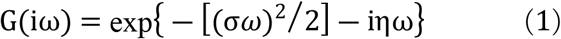

where 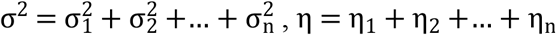 is angular frequency and i=√-1.

The real and imaginary parts of G(*iω*) are given by

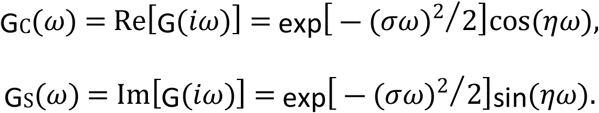

Since G(iω) belongs to the frequency domain, it can be regarded in terms of the probabilistic notions of quantum theories as a characteristic function [19]. Accordingly, the time domain counterpart of this characteristic function can be considered as a distribution.

Mathematically, the relationship between the frequency domain characteristic function F(iω) and the corresponding time domain distribution function f(t) is established by the reciprocal Fourier integrals:

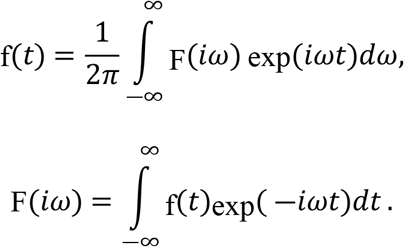

These integrals convert one of these representations into the other and vice versa. If F(iω) in the first of these integrals is expressed over an infinite frequency scale by Eq(1) i.e. F(iω) =G(iω), then f(t) is a normal distribution defined on an infinite time scale. However, we are dealing with a causal process, the development of which starts from the time instant when the triggering event activates the transient behavior of the STION. Such a condition of causality establishes specific relationships between the real, G_C_(*ω*), and imaginary, G_*S*_(*ω*), parts of G(iω). On condition that f(t)=0 at t<0, these functions mutually determine one another. Consequently, both G_*C*_(*ω*) and G_*S*_(*ω*) appear as counterparts of one and the same time domain function, and either one alone is sufficient to find the time domain counterpart, i.e. the f(t) at t≥0 [18].

Using the imaginary part, we obtain the time domain solution at t≥0 in the form of the following sine Fourier transform of G_S_(ω)

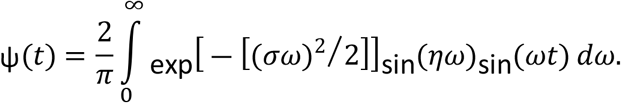

This integral has an analytical solution [21]. For t≥0

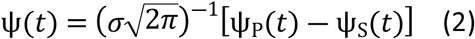

where,

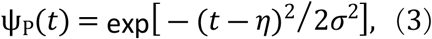

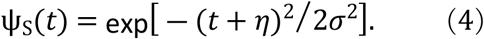

Taking *σ* = 1 and *η* = 1, these functions are illustrated in the upper panel of Fig 2 by the solid lines: ψ(t) - black, ψ_P_(t) - blue and ψ_S_(t) – red. The dotted lines are the Gaussian functions which indicate that ψ_P_(t) and ψ_S_ (t) are the fragments of the shifted normal distributions. With σ=1, the lines in the bottom panel show ψ(t) with the various values of η.

**Fig 2.**
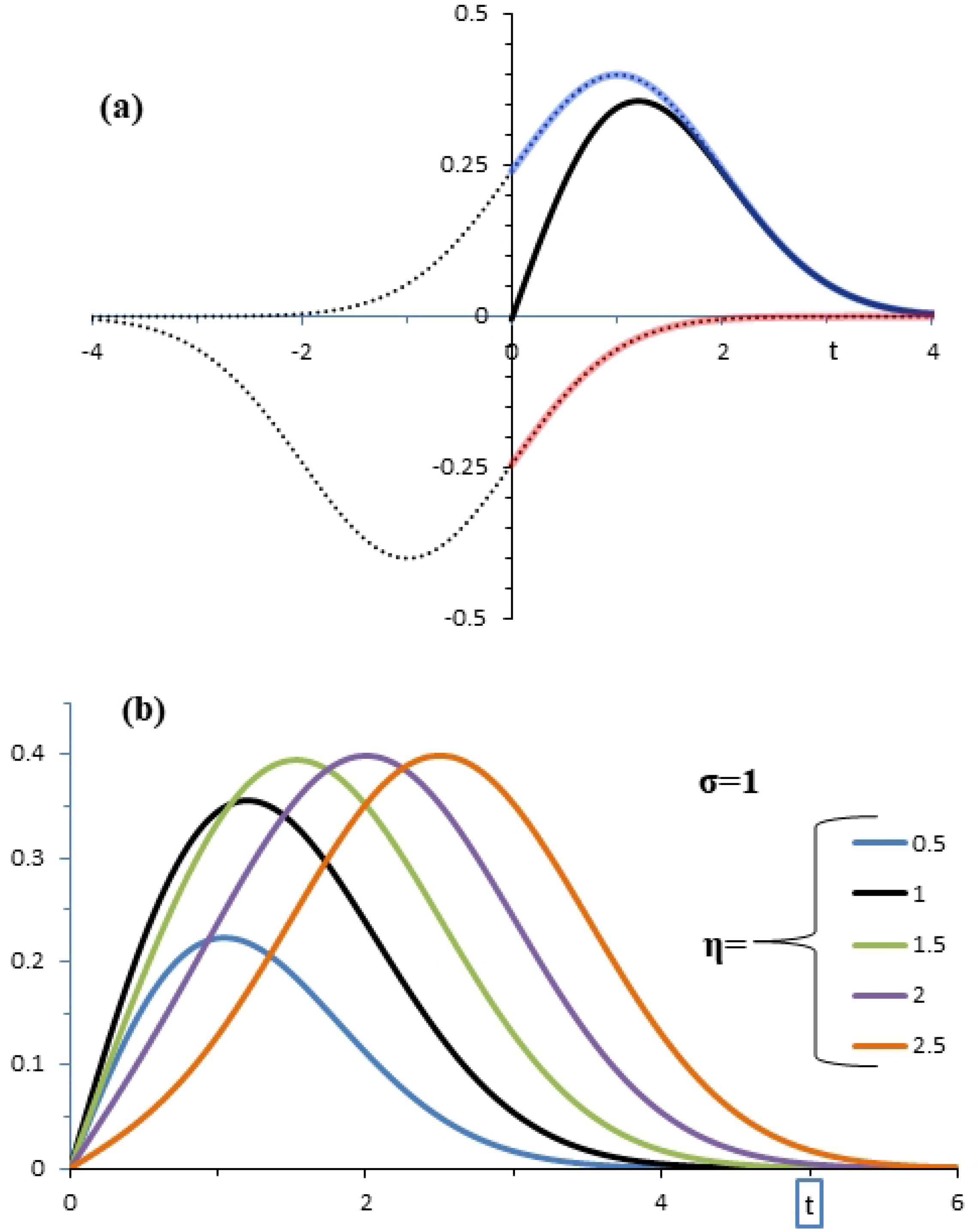
Given σ=η=1, the black, blue, and red solid lines in the top panel show ψ(t), ψ_P_(t), and ψ_S_(t), respedtively. The dotted lines are Gaussian functions which indicate that ψ_P_(t) and ψ_S_(t) are fragments of the two shifted curves of normal distributions. The bottom panel illustrates ψ(t) with σ=1 under different values of η.

Eq (2) is consistent with the *wave function* in a general form of d’Alembert’s solution to the wave equation [22]. In this context ψ(t) is the sum of a right traveling wave ψ_P_(t) and a left traveling wave ψ_S_(t). The crucial difference is that d’Alembert’s wave function is defined on an infinite time scale, *sub specie aeternitatis*, while ψ(t) is zero at t<0. This is the reason why ψ(t) is called the half-wave function (HWF).

Phenomenologically, HWF is a macroscale voltage transient which develops from t=0 as a cumulative statistical aggregate of the transient activation of multiple elementary sources. In the frequency domain this signal is defined by the complex spectrum Eq(1). We repeat this equation here in terms of the amplitude spectrum, A(ω), and the phase function φ(ω):

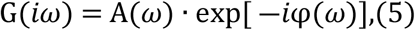

where,

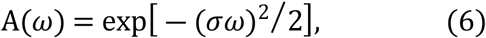

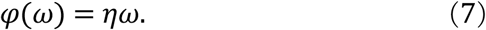

## Materials and Methods

### Subjects

Three groups of age- and sex-matched subjects have been investigated in this analysis, drawn from 2 earlier studies [12, 24]:

17 unmedicated patients with BPD, 17 patients with schizophrenia and 17 healthy controls. The BPD patients (4 males and 13 females; range = 20-44 years; mean age = 31.6) came from an ongoing program for the treatment and evaluation of BPD patients. The diagnosis was made by two independent raters (psychiatrist and psychologist), according to DSM-III-R criteria in a diagnostic interview that included the Diagnostic Interview for Borderline Patients. Patients were free of medication for at least 30 days at the time of the study [12].

The schizophrenia patients (4 males and 13 females; range = 20-44 years; mean age = 31.6) were drawn from a larger sample recruited from hospitals and community centres in Sydney. All participants had been diagnosed with schizophrenia for at least 4 years. The diagnosis was made through concordance between the case file diagnosis and diagnosis based on CIDI Section G, according to DSM-IV criteria (American Psychiatric Association, 1994) [24].

The control group included 17 matched subjects (4 males and 13 females; mean age = 34.3, sd =8.6, range = 20-47 years) [24].

Exclusion criteria for all groups were a recent history of substance abuse, past history of substance dependence, intellectual disability or other neurological disorders including epilepsy and head injury [assessed using Section M from the Composite International Diagnostic Interview (CIDI) and the Westmead Hospital Clinical Information Base questionnaire (WHCIB)]. Subjects were asked to refrain from smoking or drinking caffeine for three hours prior to the recording session. Ethics approval was obtained for the original projects from the Western Sydney Area Health Service Ethics Committee [12, 24]. Written consent was obtained from all subjects prior to testing in accordance with National Health and Medical Research Council guidelines [12, 24].

### Procedure

Subjects were seated in a sound and light attenuated room. Each subject had their eyes open and was instructed to fixate on a colored dot in the center of a screen, in order to minimize eye movements. An electrode cap [23] was used to acquire data from Fz, F3, F4, F7, F8, Cz, C3, C4, T3, T4, Pz, P3, P4, 01, and 02 scalp sites. Linked earlobes served as the reference. Horizontal eye movement potentials were recorded using two electrodes, placed 1 cm lateral to the outer canthus of each eye. Vertical eye movement potentials were recorded using two electrodes placed on the middle of the supraorbital and infraorbital regions of the left eye. All electrode impedances were less than or equal to 5 kΩ. A 32-channel continuous acquisition system with DC amplifiers was employed. EEG and EOG channels had a range of ±13.75mV and a resolution of 0.42 μV.

ERP data were collected according to a standard auditory “oddball” paradigm. Stereo headphones conveyed regular tones of 1000 Hz at an interval of 1.3 seconds to both ears. Eighty-five percent of these were 1000 Hz tones which the subjects were instructed not to respond to (task irrelevant). The other 15% were target (task relevant) tones of 1500 Hz. These high tones (targets) were intermixed with the lower (background) tones in a pseudorandom sequence, with the constraints that two target tones were never presented in succession, and the number of background tones between targets was always an odd number between 1 and 11 inclusive. Total tone duration was 60 ms, including a 10 ms rise time and 10 ms fall time.

The subjects were instructed to respond to target tones by pressing two reaction-time buttons, as fast and accurately as possible, with the middle finger of each hand (to counterbalance motor effects). Speed and accuracy were emphasized equally in the task instructions. All tones were presented at 60 dB above the subject’s auditory threshold (determined prior to recording).

The experiments for each subject consisted of a 4 min session during which the stimuli application times were recorded simultaneously and continuously with the 32-channel EEG and 2-channel electro-oculogram (EOG).

A low pass filter was applied to the analogue voltages prior to digitization. The cutoff of this filter was 50 Hz, with the attenuation being 40 dB per decade above 50 Hz. In addition, a 50 Hz notch filter was applied to eliminate 50 Hz AC mains power supply interference.

Filtered voltages were continuously digitized at 250 Hz and digitally stored with the markers of the instants of stimuli applications.

### Fragmentary decomposition

There is general agreement that EEG and ERP are complex signals composed from multiple components produced by activation of various ensembles of cortical cells. The components of EEG and ERP signals are usually defined differently. With the EEG, most of the methods consider a signal as a composite of several oscillatory components the most important of which are the delta, theta, alpha and beta band rhythms. In contrast, ERP components are defined in terms of peaking waveforms with characteristic polarities and latency ranges [1].

An appealing feature of the probabilistic methodology introduced above is that it provides a universal description of EEG and ERP components in the form of Eq 2, i.e. the HWF. The probabilistic background of this concept has been developed in a series of previous studies, and empirically supported by the time-frequency analysis of a number of biomedical signals: the EEG [24], event related potentials (ERP) [25], electrocardiogram [26] and eye-blink electromyogram [27].

Conceptually, the HWF corresponds to a conventional component definition, according to which a peaking waveform may be considered as a major descriptive element of the electrophysiological signal.

Using a model component in the form of HWF, the model of an EEG segment with “N” components has the form of the following sum of weighted HWFs

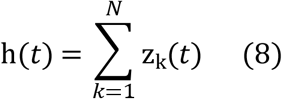

where z_k_(t) = g_*k*_ · ψ_*k*_(*t* – *τ*_*k*–1_).

The index “k” in this formula labels different HWFs with corresponding σ_k_ and η_k_ parameters. g_k_ is the weighting coefficient and τ_k_ is the time instant from which the development of the corresponding component starts.

We identify this model (i.e. the estimation of the parameters of each term in the right hand side of Eq (8)) using the method of fragmentary decomposition (FD) with some new features [27, 28].

The EEG signal for FD is given in a digital form of the time series **V** = 〈v_0_,…,v_*i*_,…,v_*I*_〉, where v_i_ = v(t_i_) are the EEG samples at discrete regular time points t_*i*_ = *i*Δ (Δ is the sampling interval). The value of Δ is chosen to exceed the Nyquist rate, being the lowest sampling rate that is necessary for accurate restoration of the signal v(t).

One class of techniques, considered as a straightforward fragmentary decomposition, defines the empirical counterpart of HWF as the EEG fragment over which the peaking waveforms of the signal are developed [25]. The fragments are separated by the segmentation points, defined as zero-crossings and points of the global and local minima in the time course of |v^(t)^|.

More particularly, if,

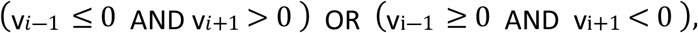

then τ_k_ =t_i_ is qualified as a zero-crossing (segmentation point), where index *“k”* is the number of the segmentation point. The point τ_k_ =t_i_ (k is the number of the segmentation point) specifying a global or local minimum of v(t) if |v_i-1_|≥|v_i_|≤|v_i+1_|.

By ordering the segmentation points as successive time instants, the sequence of the segmentation points τ_0_,…, τ_k_,…,τ_P_ is formed. Taking a segment of the length T_k_ = τ_k_-τ_k-1_ between the segmentation points τ_k-1_ and τ_k_, the empirical half wave (EHW) is defined as 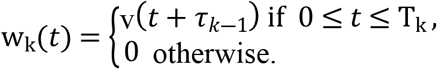

In the interval from 0 to T_k_ this function reproduces the signal fragment between the adjacent segmentation points τ_k-1_ and τ_k_.

Given P+1 segmentation points in the interval from τ_0_ to τ_P_,

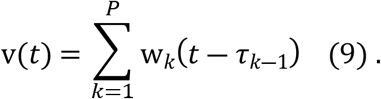

The comparison of Eq(8) and Eq(9) signifies w_k_(t) as an empirical counterpart of z_k_(t).

The limitation of the straightforward fragmentary decomposition is that it does not consider the possibility of component overlap. To avoid the influence of component overlap on modelling accuracy, the method of high-resolution FD (HRFD) has been developed [28]. HRFD is organized as a recursion procedure performed in successive steps from the first term to the last term in Eq (9).

The procedure involves P successive steps each of which estimates a particular z_k_(t) from Eq (8). At the m*th* step (m<N), the Eq (8) is made up of “m” terms

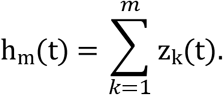

The development of z_m_(t) starts from the segmentation point τ_m-1_ which corresponds to the sample X = *τ*_*m*-1_/Δ. In straightforward fragmentary decomposition, the estimation of the segmentation point is supported by the analysis of the time course of the original signal v(t). This means that z_m_(t) serves as a model of the signal on the internal from τ_m-1_ to τ_m_. Actually, the development of z_m_(t) may continue at t> τ_m_. This produces an overlap of z_m_(t) with the next term, i.e. z_m+1_(t). The component overlap is quantified by the residual, r_m_(*t*) =v(*t*) - h_m_(*t*).

This residual serves for the estimation of the next segmentation point τ_m_ which is defined as a zero crossing or minimum in the time course of |r_*m*_(*t*)| at t> τ_m-1_.

Taking the signal segment of the length L_k_ = τ_m_-τ_m-1_ between the segmentation points τ_m-1_ and τ_m_, the overlap corrected EHW is defined as

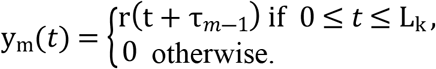

Since the following transformations of y_m_(t) to the frequency domain are universal the “m” subscript is omitted and y(t) denotes the overlap corrected EHW defined on the interval [0,L].

The frequency domain counterpart of y(t) is defined by the exponential Fourier transform

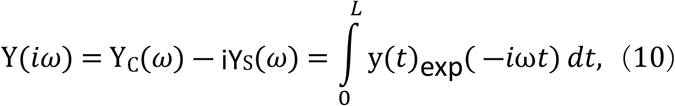

where ω=2πf, f is frequency and i=√-1.

Since y(t) is an empirical entity, the calculations of such trigonometric integrals necessitate numerical algorithms.

The numerical method supporting fragmentary decomposition is the similar basis function (SBF) algorithm [29], a novel version of Filon-type methods which provides maximum precision in the estimation of trigonometric integrals. The SBF algorithm deals with the continuous Fourier spectrum instead of a discrete spectrum defined by the discrete Fourier transform.

This allows employment of a logarithmic frequency scale for calculations and display of frequency characteristics. Another important aspect of this innovation is that the SBF algorithm overcomes well-known difficulties of short-time period spectral analysis and is applicable to signal segments of arbitrary length without windowing and zero-padding. This is especially important for algorithms dealing with EHWs, the lengths of which are variable.

To estimate complex Y(iω) using real numbers, the numerical procedures of the SBF algorithm are separately applied to the real and imaginary parts of the complex spectrum defined by Eq(10). These are:

i. the finite cosine Fourier transformation

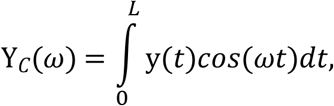

and
ii. the finite sine Fourier transformation

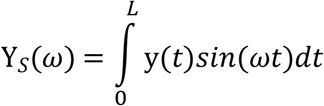

where ω=2πf is the angular velocity, and f is the frequency.

The corresponding amplitude spectrum and phase function are defined as:

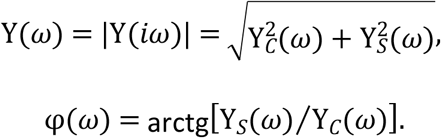

These are continuous functions of angular frequency, the values of which can be calculated by the SBF algorithm for arbitrary sets of points. A logarithmic frequency scale is employed for calculations of amplitude spectra and a natural frequency scale for calculations of phase functions.

In the case of the logarithmic scale, frequency characteristics are calculated for angular frequencies ω_i_=ω_0_C^i-1^ (i=1,…, N), where ω_0_ is initial angular frequency and C>1 is the parameter which defines the sampling rate. The selection of this parameter is supported by the formula C=exp(ln10/N_D_), where N_D_ is the number of samples per decade.

The amplitude spectrum and frequency are normalized in order to express these entities in dimensionless units.

The normalized amplitude spectrum is defined as

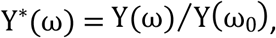

where Y(ω) is the computed amplitude spectrum and ω_0_ is a sufficiently small value of angular frequency selected to satisfy the condition: Y(ω_0_) ≈ Y(0). The blue line at the middle panel of Fig 3 illustrates a typical normalized amplitude spectrum. It was calculated from the EHW depicted at the top of the figure. The red line shows the fit of the theoretical amplitude spectrum (6) to the empirical Y*(ω).

**Fig 3.**
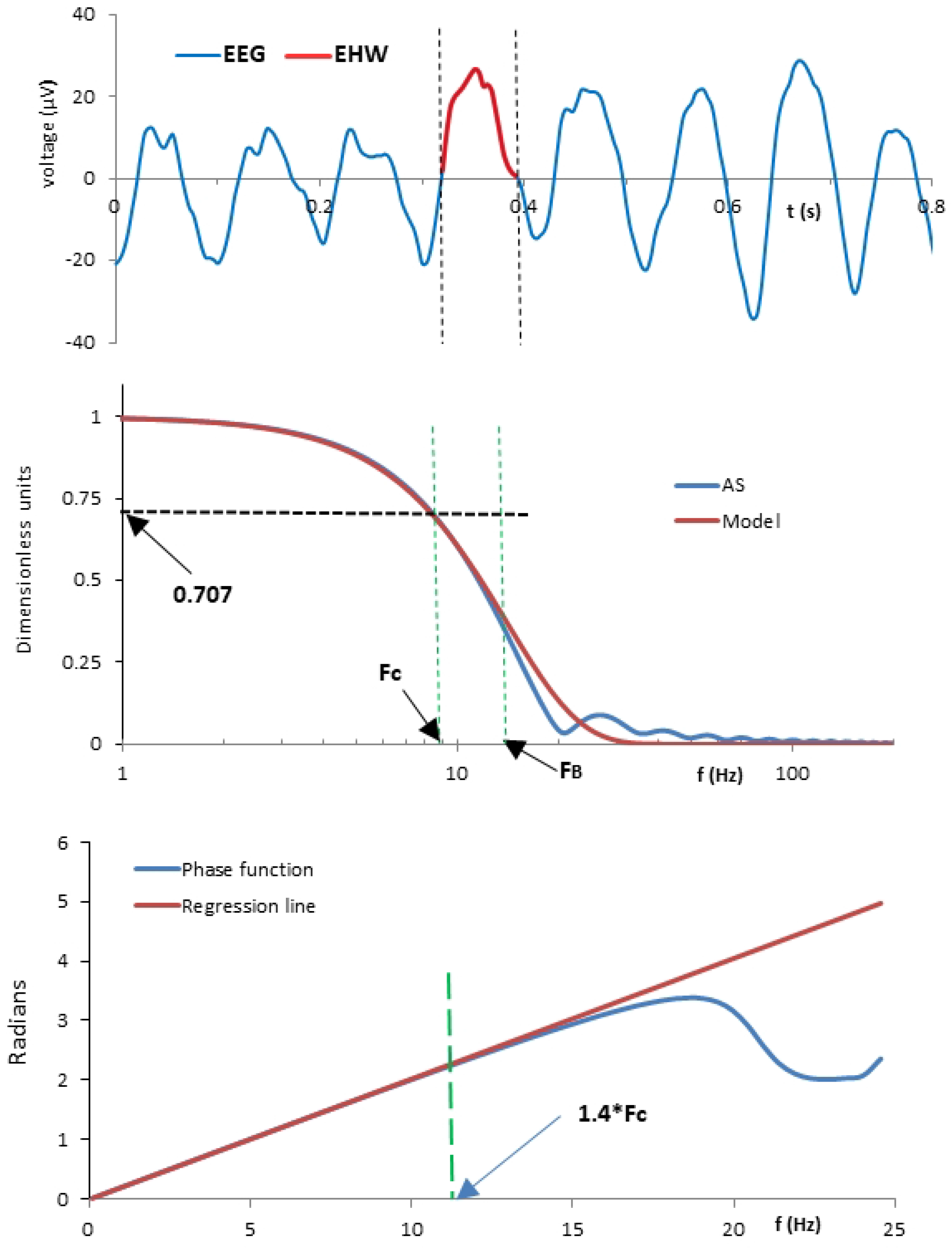
Top panel: the blue line shows 800 ms segment of typical EEG, the vertical dotted lines delineate the EHW. The blue lines in the middle and bottom panels show the normalized amplitude spectrum and the phase function calculated from selected EHW. The red line in the middle panel shows the Gaussian template at position defined by the best fit. The red line in the bottom panel is a regression line which illustrates typical linearity of the phase function in the frequency range from F_0_ to 1.4·F_C_.

An important factor supporting the fitting procedure is that Y*(ω) can be considered as the frequency response of a low pass filter, the conventional parameter of which is the cut-off frequency FC. At this frequency the attenuation of the amplitude spectrum drops by 3dB, i.e. 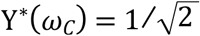, where ω_C_=2πF_C_.

The cut-off frequency serves as a basis unit, the use of which defines the relative frequency as γ=ω/ω_C_.

Using the dimensionless amplitude and frequency, the empirical amplitude spectrum is defined in relative units as Z(*γ*) = Y*(*ω_C_γ*). The corresponding model is the Gaussian spectrum

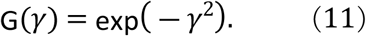

Note that Z(*γ*) = G(*γ*) at *γ* = 1, the relative frequency which corresponds to f=F_C_.

In most cases combining Z(γ) and G(γ) at γ=1 provides a close agreement between these entities. In a systematic manner the comparison is performed more fully by a fitting procedure, the goal of which is finding of the best match of analytical G(γ) to empirical Z(γ).

The fitting uses G(γ) as a template which is moving throughout the abscissa scale. During this procedure the template samples remain unchanged while their locations on the abscissa scale (dimensionless γ) are jointly shifted by the multiplication of relative frequencies by an appropriate constant.

Starting from the frequency F_0_=ω_0_/2π, the accuracy of the fit is evaluated by the value of the mean square error between Z(γ) and G(γ). The position of the best fit defines the boundary frequency F_B_. Numerous trials with various EHWs revealed that typically the template from the best fit virtually coincides with the amplitude spectrum in the range of standard frequencies from 0 to 1. At γ>1 the errors increase with increases in frequency.

The larger that F_B_ is in comparison with F_0_, the more accurate the Gaussian model of the amplitude spectrum. For assessing a goodness of fit, the dimensionless extension ratio ε=F_B_/F_C_ is used. The fit in the middle panel of Fig 3 gives a visual idea of how the F_C_ and FB are related.

Calculations of σ are followed by the estimation of phase functions. It was found that the initial part of the phase function φ(ω) shows consistency with a simple linear model

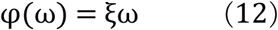

where ξ is a parameter.

Numerous calculated linear fits indicate that deviation from linearity can be neglected over the frequency range from 0 to 1.4·F_C_. A typical result is illustrated in the bottom panel of Fig 3. Thus, the estimation of η is reduced to the calculation of the linear regression line using φ(f) samples from f=f_0_ to f=1.4·F_C_. The slope of the regression line ξ serves as the estimate of the parameter η.

Given particular w_k_(t) from Eq(8), the estimated σ and η define the corresponding ψ_k_(t) term. Using these data, the weighting coefficients are derived from the following interpolation conditions separately applied to each pair of EHW and HWF:

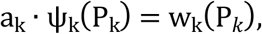

where P_k_ is the peak latency of the *k*th EHW. Consequently,

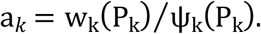

Thus, the peak latencies and amplitudes of the model are equalized to the peak latencies and amplitudes of EHW.

### Single trial analysis

The HRFD supports an original method of single trial analysis of ERPs. The procedure starts from the extraction of EEG segments time-locked to the target stimuli. Given a particular EEG channel with a digitally stored data set, for each of the first 40 target stimuli, standard 2s EEG segments were extracted, from 1s pre-stimulus to 1s post-stimulus. The segments were digitally filtered (moving window averaging) to remove irrelevant low (<0.5 Hz) and high (>50 Hz) frequency components. For each channel these procedures provided a time series with 500 samples v[nT], where n takes values −250,..,0,..,249 (n=0 is the time of the target stimulus onset) and T=0.004 s.

The application of the HRFD provides a model of a single trial in the form of Eq 8. The parameters σ_i_ and η_i_ of each identified HWF are transformed to conventional parameters: A_i_=0.356σ_i_ - peak amplitude and L=t_i_+1.2η_i_ - peak latency. Physically, 1.2η is the rise time, i.e. the time interval during which the component increases from zero to its maximum value. This measure becomes available due to the shape estimate (parameter η) provided by the FD technique.

An extended system of parameter windows has been developed for identification of conventional late ERP components N1(00), P2(00), N2(00), P3a and P3b. Given L - peak latency, A - peak amplitude and η – shape constant as the major parameters, the identified HWF is qualified as a meaningful ERP component if it satisfies the conditions specified by the following windows:

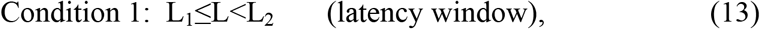

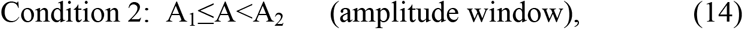

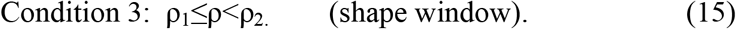

The L_1_ and L_2_ (ms) parameters of the latency windows were as follows: P50 – 20 and 75, N1 - 80 and 120, P2 - 160 and 220, N2 - 180 and 235, P3a - 240 and 299, P3b - 300 and 360. The parameters of the shape windows for all components were: ρ_1_=8 ms and ρ_2_=50 ms. The A_1_ and A_2_ (μV) parameters of the amplitude windows were −45 and −2 for negative and 2 and 45 for positive components, respectively.

### Averaging procedures

Contemporary understanding of endogenous potentials is mostly based on the visual examination and quantitative analysis of average ERPs.

The traditional method of averaging assumes that single trial recordings consist of identical time-locked ERPs and random constituents that are not time-locked to the event. However, it is widely accepted that this assumption is an oversimplification which discounts the reality that ERP composition is made up of multiple components, each of which is subject to trial-to-trial variability which may be governed by different factors. Elucidation of the nature of these components requires explicit analysis of various ensembles of identified single trial ERP components.

Our methodology of HRFD identifies ERP components in each single trial and creates a model of single trial ERP in the form of Eq 8. This solution provides a basis for a novel method of averaging which we call selective component averaging (SCA). The averaging is applied to identified HWFs which satisfy conditions 1-3 for the component of interest. The SCA of selected components is defined by the sum

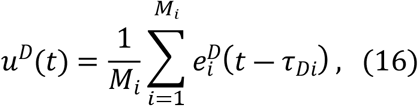

where the symbol “D” stands for the name of the component, M_i_ is the number of selected HWFs and τ_Di_ is the time instant from which the HWF starts. The sum of models defined by Eq16 is called a synthetic average. For example, synthetic average of the late positive complex consisting of the P3a and P3b has the form,

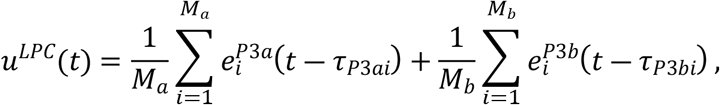

where *M_a_* and *M_b_*, are the numbers of *P*3_*a*_ and *P*3_*b*_, components delivered through SCA.

SCA improves the accuracy of average waveforms because it selects the trials with identified (i.e. meaningful) components and ignores trials with missing components.

To account for missing responses, we introduce a novel parameter called an elicitation rate (ER). This parameter takes into account an actual number A of the trials in which the component was defined: A=T-M where T is the total number of single trials and M is the number of the trials with missing components. The ER is defined as P_E_ = A/T. This parameter is the probability of the elicitation of a defined component in a single trial.

### Statistical analyses

#### Kolmogorov Smirnov test

The procedures of parameter estimation described above define the frequency range of the best fit of the theoretical amplitude spectrum to the empirical amplitude spectrum of EHW from F_0_ to FB. Comparison of different fits is enabled by the dimensionless extension ratio ε = F_B_/F_C_. Given the samples of ε in the form of two different ensembles, 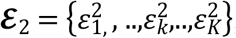 and 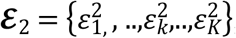, the Kolmogorov-Smirnov one and two sample tests are used in order to decide whether 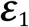 and 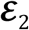 are produced by the same or different distributions. Each of the data sets 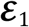 and 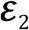 is converted to a cumulative frequency distribution. The test is based on evaluation of the maximum vertical deviation D between the cumulative frequency distributions. The null hypothesis that the two distributions are the same is rejected if the value of D exceeds the critical value defined by the tables of D statistics.

## Results

Our theory proposes that scalp potentials are produced by collective behaviour of multiple LCGs, the general properties of which are defined in terms of the STION and HWF. The results we report are dedicated to the micro-scale and macro-scale phenomena supported by these concepts.

### Probabilistic treatment of micro-scale phenomena

According to our theory, the central role in microscale events belongs to the cellular membranes which control the transport of the ions from the intracellular to extracellular space and vice versa. The key idea we adapt from previous work [5], is to describe these transport processes in terms of a birth and death process (BDP). Physically, a particle moving out of a cell would constitute a ‘death’ for the inside of the cell and a ‘birth’ for the extracellular space.

The state of the STION is defined by the net number of positive and negative ions released from the cells of the LCG. It is measured at time t by the integer-valued time-dependent random variable **X**(t) which we consider as a Markov process [20]. The main assumption of the Markov process is that during a sufficiently small element of time, Δ, the probability of the change of **X**(t) by more than one particle is negligibly small:

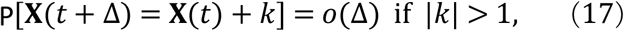

where P denotes probability and k is an integer.

A remarkable property of the Markov process seen in this equation is that the future state of the population (i.e. **X**(t+Δ)), depends only on its present state **X**(t), and not on how that state was reached. Therefore, we deal with a Markov chain, a stochastic model describing a sequence of possible events in which the probability of each event depends only on the state of the previous event.

A particle system can change its state only through transition to the nearest neighbourhood. An increase of the population size by a unit represents a birth, **X**(t+Δ)=**X**(t)+1, whereas decrease by a unit represents a death, **X**(t+Δ)=**X**(t)-1. Wide classes of BDPs with constant transition probabilities are associated with stationary processes. However, the complex dynamics of EEG signals indicates both highly irregular and non-stationary behaviour of the underlying particle systems. We use a nonhomogeneous BDP, as the most suitable mathematical tool, in which the birth and death rates may be any specified functions of the time t [30]. The probabilities of the changes of the population size for a nonhomogeneous BDP are:

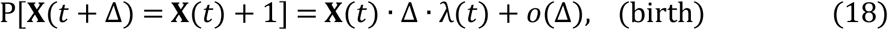

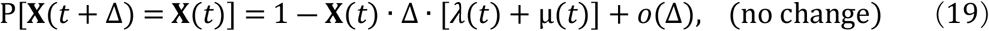

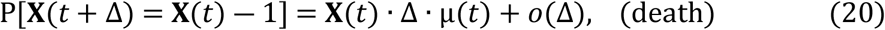

where λ(t) and μ(t) are the time dependent birth and death rates, respectively.

Let us choose Δ in compliance with Eq(17) and denote by x_i_ = X(*t_i_*) the state of a STION at the time ti. Using these terms, we can represent the time evolution of **X**(t) as a succession of discrete states x_i_. Permitted states of the particle population at the time t_*i*+1_ are: xover 80ms time interval were_i+1_=x_i_+1 (birth), x_i+1_=x_i_ (unchanged size), or x_i+1_=x_i_-1 (death). The probabilities of the corresponding inter-state transitions are:

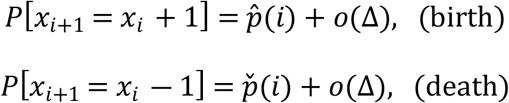

where 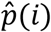 and 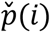 denote the time dependent transition probabilities for the birth and death, respectively. From Eq(18) and Eq(20) we get:

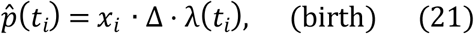

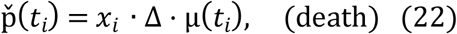

These relationships are adequate tools for the analysis of particle movements (i.e. microscale events).

### Bridging micro and macro-scale phenomena

To proceed from our treatment of micro-scale events to description of macroscale processes we need to express cumulative effects of micro-scale events in deterministic terms. An important point for the solution of this problem is that a deterministic model is a special case of a stochastic model, in the sense that it yields results which hold with probability one. Consequently, the expression for a mean population size from the model of stochastic BDP is the same as that for the population size obtained from a deterministic kinetic model.

In view of this correspondence, we consider the mass effect produced by the STION in terms of a mean size described by the function ψ(t). We assume that ψ(t) develops from the time instant t=0 when some triggering event transfers the LCG from the resting state to the transient conditions. The expected trajectory of the mean size of the particle population at t≥0 is [30]

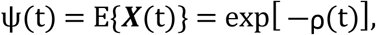

where E {} denotes expected value and

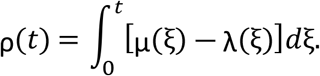

Thus,

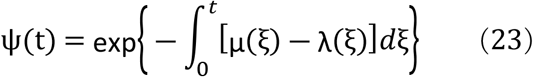

According to the theory of field potentials within the frequency range of physiological interest, the extracellular space may be regarded with reasonable accuracy as a resistive medium [15]. This means that the tissue that lies between the STION and the ground acts as a resistive volume conductor. Therefore, the temporal evolution of the global scale transient potential v(t) is proportional to the expected population size of the STION. This means that v(t)=k·ψ(t), where *k* is the weighting coefficient.

### Primary and secondary populations of ions

The two terms from which the HWF is composed are considered as the products of two sub-populations of particles: the *primary particle population* associated with ψ_P_(t) and the *secondary particle population* associated with ψ_S_(t).

We deduce the parameters η and σ of these populations from the EEG dynamics on the macroscopic scale. The integral in Eq (23) allows us to estimate these macroscale measures using λ(t) and μ(t) variables which govern the microscopic scale processes.

The BDP with the birth λ_P_(*t*) and death μ_P_(*t*) rates, serves as a model of the primary particle population. Accordingly, ψ(t) in Eq (23) is replaced by ψ_P_(t) from Eq(3), which gives

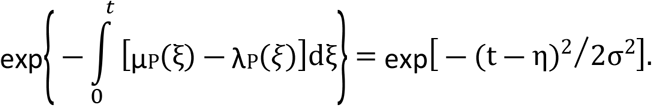

Consequently,

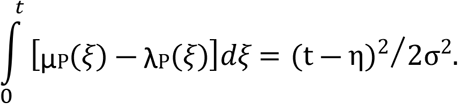

The solution of this equation is a simple matter that demonstrates

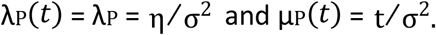

Following the same reasoning as above, the secondary particle population is expressed in terms of ψ_S_(t), and the birth and death rates denoted by λ_S_(t) and μ_S_(t).

The replacement of ψ(t) in Eq (23) by ψ_S_(t) shows

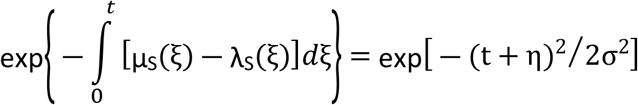

Consequently,

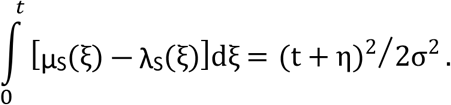

It follows from the solution of this equation that

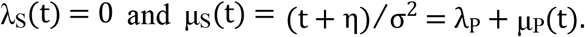

As in [6], we summarize these relationships in the form of the following probabilistic rules which define the birth and death rates for the primary and secondary particle populations.

#### Rule 1.

*After activation at t=t_0_ by the triggering event the transient behavior of the primary particle population develops as a non-homogenous BDP with a constant rate of birth*

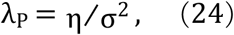

*and time-dependent rate of death*

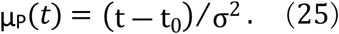

#### Rule 2.

*After activation at t=t_0_ by the triggering event the transient behavior of the secondary particle population develops as a non-homogenous death process with a timedependent rate of death*

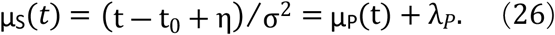

With respect to resting conditions, it is essential that ion transport is balanced for both cations and anions. This implies that, in the resting state, the sizes of the primary and secondary particle populations vary around the mean values. These processes can be considered as simple BDPs with constant rates of birth and death. An important point which defines these parameters, is that Equations (24)-(26) contain the time-dependent term μ_P_(t) and time independent term λ_P_. This means that λ_P_ participates in both the resting and transient conditions. It is a unique parameter which controls both the birth and death processes during the resting conditions. Accordingly, the BDPs in the primary and secondary particle populations are governed, under resting conditions, by the birth and death rates equal to η/σ^2^. The corresponding transition probabilities are:

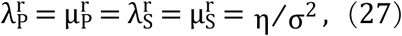

where 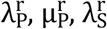 and 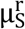 denote the resting state rates of birth and death for the primary and secondary particle populations.

Numerical simulations are unique tools for reconstruction of the time courses of particle populations in different trials. To deal with particle numbers, the numerical counterpart of equation (2) is presented in the form

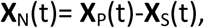

where **X**_N_(t) is the number of particles corresponding to the net charge, while **X**_P_(t) and **X**_S_(t) are the numbers of particles in the primary and secondary populations, respectively.

In the following general cases the subscript is omitted for brevity. Thus, x_i_ denotes the number of particles at time instant t_i_, that is, x_i_ **X** (t_i_).

Particle populations are considered at time instants t_i_=iΔ (i=0, 1,..), where t_0_ is the time at which the EEG transient starts. With reference to (21) and (22) the following formulas of transition probabilities correspond to the transient conditions.

Primary particle population:

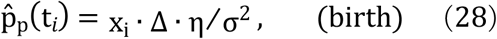

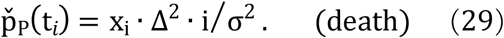

Secondary particle population:

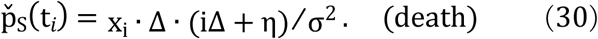

According to Eq(27) the birth and death processes in the primary and secondary particle populations are governed under resting conditions by the birth and death rates equal to η/σ^2^. The corresponding transition probability is

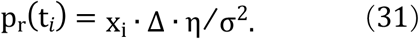

Eqs (28)-(30) provide a means for creating numerical reconstructions of the time evolution of particle populations during transient and resting conditions. A general procedure is organized as the succession of standard steps dealing in sequential order with the time intervals [t_i_, t_i+1_] for i taking values from -M to N-1. The time interval from t_-M_ to to corresponds to resting conditions. At t=t_0_ the resting state is switched to the transient conditions.

The simulations illustrated in Fig 4 extend over the time interval from −10 to 70 ms with t=0 corresponding to the instant at which the transient condition starts. As an initial condition, an equal size N_0_=50 was prescribed for both populations with parameters σ=13.3 ms and η=26.2ms.

**Fig 4.**
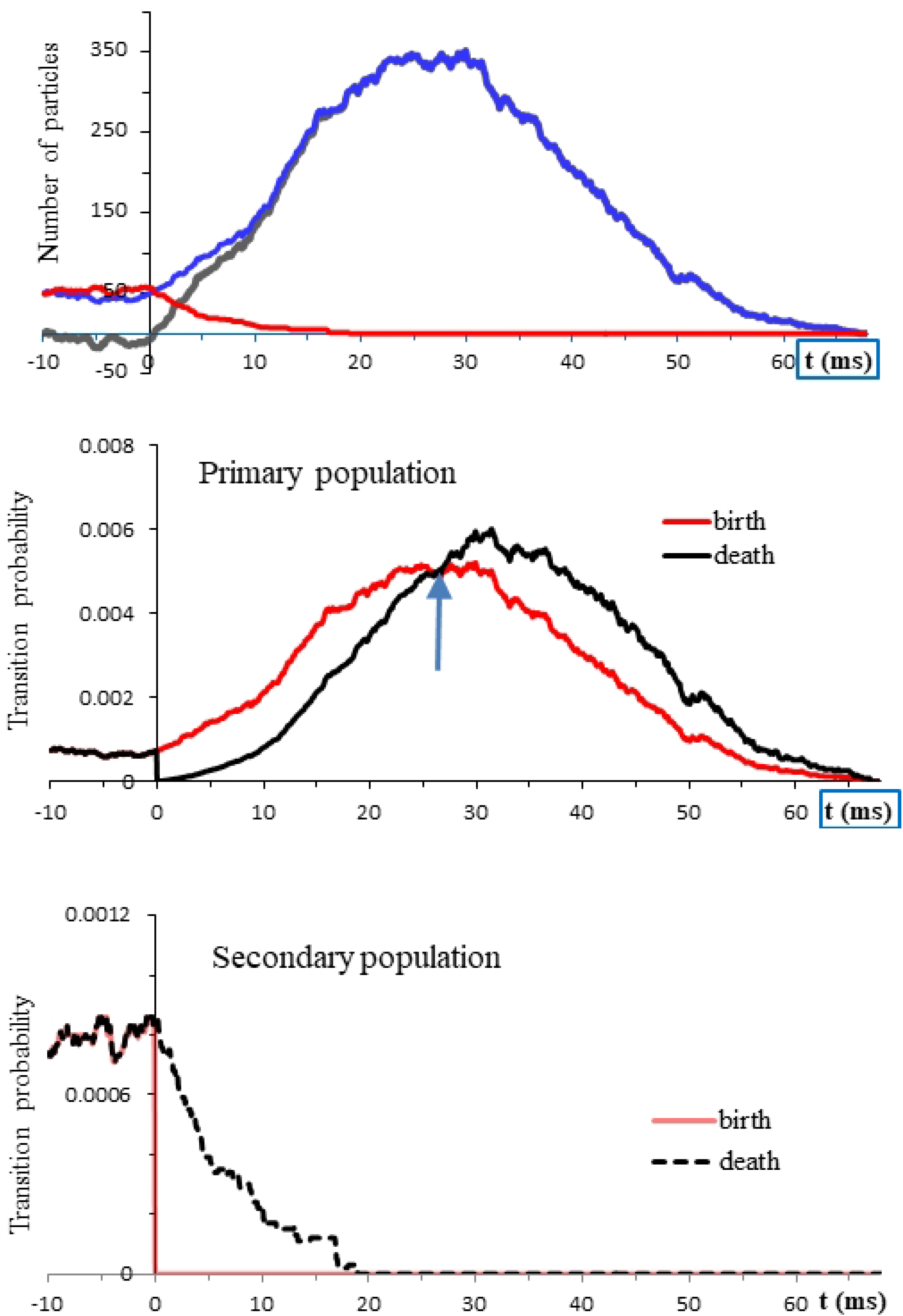
Numerical reconstructions of the temporal evolution of particle populations in a typical trial. The resting conditions computed from −10 ms are turned at t=0 to the transient conditions. The blue, red, and black lines in the top panel show the functions *X*_P_(t), *X*_S_(t) and *X*_N_(t). The time courses of the underlying transition probabilities are shown in the middle and bottom panels.

Using defined parameters, it was important to satisfy the condition of Eq (1) through choice of a sufficiently small time-interval Δ, during which the expected change of the population size by more than one particle is negligibly small. Using Monte Carlo simulations for estimation of the numbers of particles crossing the membrane, different values of Δ were tested. In this way the value Δ=0.0001ms was selected for the following simulations. The corresponding segmentation points over 80ms time interval were t_i_=i·Δ with *i* taking values from 0 to 800,000.

The transition from resting to transient condition was simulated as the change of the resting state transition probabilities, defined by Eq 21, to the transient state transition probabilities in Eqs (18-20). The change occurs in a “smooth” fashion. This means that the sizes of the primary and secondary particle populations developed under resting conditions serve as initial conditions for the transient regimes.

During resting conditions (interval from −10ms to t=0) the transport of particles between the primary and secondary populations is balanced. Particle numbers fluctuate over the mean value equal to N_0_. The manner in which the transition from resting to transient conditions contributes to a rapid change of net charge is due almost entirely to the change of birth and death rates in the primary particle population. In the general case, the size of the primary population is governed by the complex interplay of birth and death transition probabilities. The onset of transient conditions gives rise to both probabilities. Initially, from t=0 to the time instant indicated in the middle panel of Fig 3b by the arrow, the birth probability prevails over the death probability. At this stage, a near-tenfold increase of the size of the primary population occurs. After the peak, death probabilities take a progressively larger share. As a result, the size of the primary population declines and returns to initial conditions.

The effect of the transients in the secondary particle population on the net charge is minor and brief compared with the primary population. As shown in the bottom panel of Fig 3, at t=0 the probability of birth in the secondary population drops to zero. This means particle transfers from the inside to the outside of cells are effectively blocked.

In order to decide whether the mass effects of particle movements are sufficient to fully account for the dynamics of macroscale processes, it is necessary to compare the results of computer simulations with the theory. In agreement with Eq (8), we consider transient conditions starting at t=0. Since ψ(0)=0, we use ψ_0_=ψ_P_(0)=ψ_S_(0) as the initial condition for theoretical solutions. The corresponding initial condition for numerical simulations is **X**_P_(0)=**X**_S_(0)=N0. Defined ψ_0_ and N_0_ parameters allow us to create dimensionless functions ψ*(t)=ψ(t)/ψ_0_ and 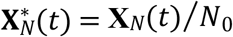. After this normalization we can directly compare the numerically calculated 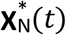 with the theoretical ψ*(t).

Taking values of σ and η parameters from the previous simulations, the samples of 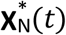 computed with N_0_ equal to10, 20, and 100 particles are illustrated by the colored lines in the top panel of Fig 5.

**Fig 5.**
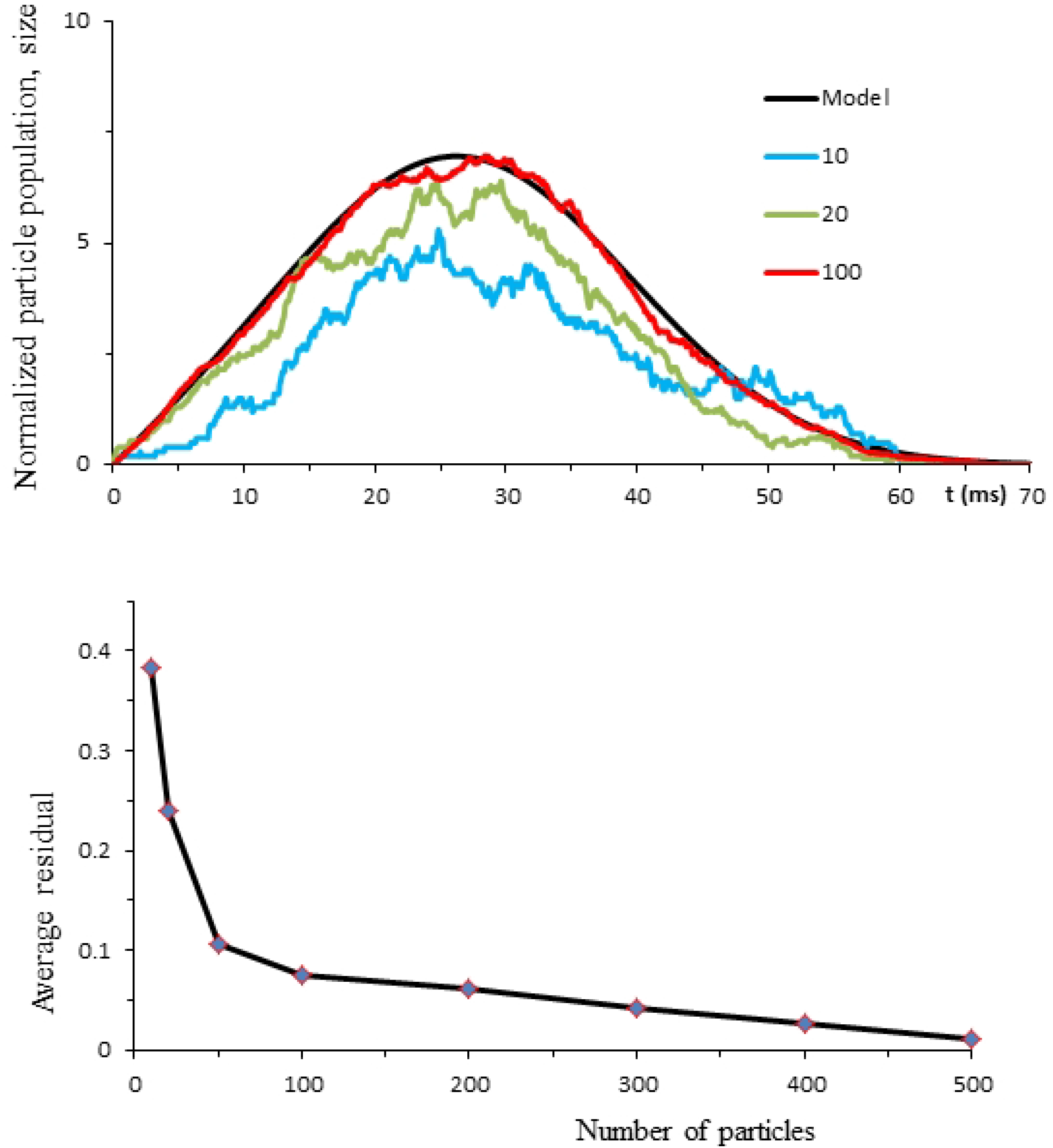
Coloured lines in the top panel show typical computer reconstructions of the temporal evolution of particle populations with different initial sizes of 10, 20, and 100 particles. The black line is the theoretical solution η^*^(t). Data points in the bottom panel show average residuals between the theoretical and numerical solutions calculated for the particle populations with different initial sizes.

It is evident that an increase in the number of particles brings single trial samples of ψ^*^(t) to a better proximity with the theoretical model. Thus, the role of deterministic factors in statistical samples of 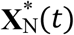 increases with an increase in the number of particles involved.

To emphasize this tendency, we have estimated absolute values of residuals between ψ*(t) and 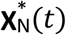. Given N_0_, 20 single trials were calculated. In each trial the residuals were estimated at 100 equidistant points. From these measurements the average residuals were calculated. The points in the bottom panel of Fig 5 show the dependency of average residuals on the number of particles N_0_.

The increase of particle numbers to several hundred makes single trials identical to the theoretical solution. This provides convincing evidence that, under transient conditions, particle behavior in both primary and secondary particle populations develops as a specific amalgamation of deterministic and stochastic processes. In an earlier study of mass potentials this phenomenon was called “transient deterministic chaos” [5].

### Transient deterministic chaos

Research into the phenomena of chaos is progressing steadily, extending to a diverse range of applications. No universal definition of the system producing chaos has been accepted, leaving us in a situation of evolving hypotheses and concepts. The necessary conditions for defining chaotic phenomena are non-linearity of the system generating chaos and its sensitivity to initial conditions.

In terms of the specifics of biological systems, in most cases the equations for the sources of deterministic chaos are unknown [31]. Accordingly, the identification of chaotic behavior is usually based on analysis of empirical time series. The most popular methods for characterizing chaotic behaviors are the Lyapunov exponents and the Kolmogorov-Sinai entropy. These are essentially indirect methods which do not define equations but diagnose whether the process in question is chaotic or non-chaotic. This limitation applies equally to other diagnostic methods, amongst which are the Poincare map, correlation dimensions, fractal dimensions, and attractor reconstructions from time series.

Our approach actually does not need such indirect methodbecause the theory provides complete models of the HWF generation on both macro- and micro-scales. The two accepted major criteria which allow us to qualify the system producing these processes as chaotic are as follows:

1. The non-linearity of the system producing HWF.
2. Sensitive dependence of the system behavior on initial conditions.

The nonlinearity of the system is evident on the macroscopic scale where the system producing the HWF is expressed by the following system of nonlinear differential equations:

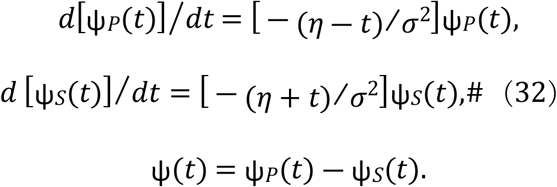

We may qualify the system as a non-autonomous nonlinear system, that is, a system driven by time-varying processes.

The strong dependency of the chaotic processes in question on initial conditions is seen from the computer simulations illustrated in Fig 6.

**Fig 6.**
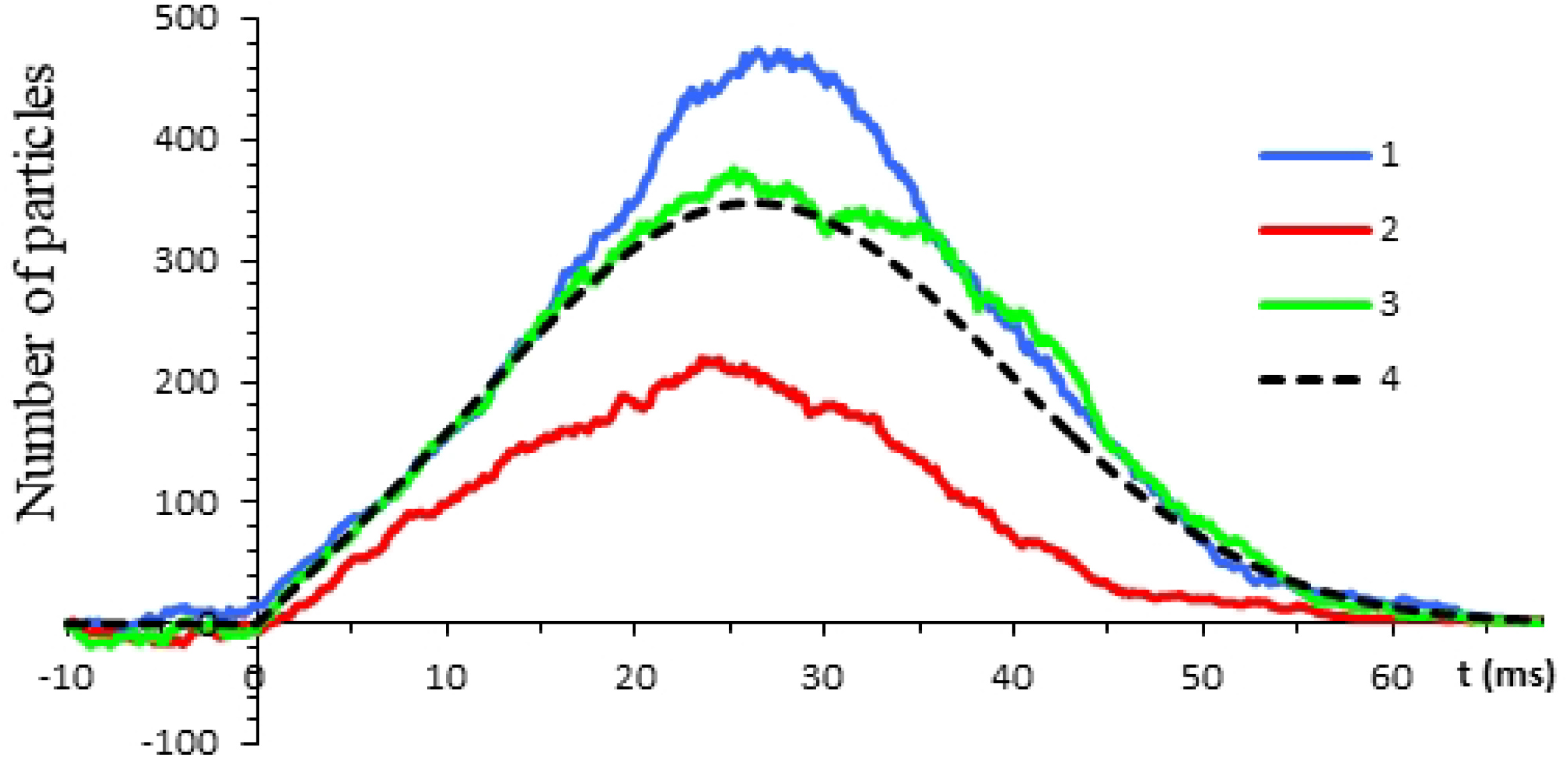
Using the same initial conditions as the simulations in Fig 4, illustrates the sensitive dependence of **X**_N_(t) realizations in different trials (colored lines) on initial conditions. The dotted line is a theoretical solution.

The parameters are the same as in the simulations shown in Fig 3. The simulations started from 10ms before the arrival of the triggering event. At that instant both P and S populations contain 50 particles. Because in the initial 10 ms interval the sizes of the P and S populations were changing randomly as simple BDPs, the initial size of the whole population was slightly different in different trials. As the results of the simulations indicate, these minor differences in the initial conditions led to significantly different future behavior in the realizations of **X**_N_(t).

Another important feature of chaotic processes is short-term predictability. It is clear from the simulation results in Fig 6 that, despite substantial differences in the trajectories of **X**_N_(t) realizations, it is possible to give rough estimates of some parameters, for example the times at which the trajectories reach their maximums. We may also claim that the feature of predictability is also incorporated in the time courses of **X**_N_(t) realizations because the averages of these trajectories from different trials converge to the analytical solutions.

### Macro-scale phenomena: EEG, ERP, P3a, P3b

The analysis of various EEG and ERP recordings in both the patient and control groups and the results of many numerical experiments provide ample evidence that half-wave function ψ(t), emerging as the macroscopic scale effect from synchronized chaotic ion movements on the microscopic scale, can be regarded as a universal building block from which these signals are composed. The corresponding empirically testable models are expressed by Eq(8).

Typical results addressed to an ongoing EEG are illustrated in Fig 7.

**Fig 7.**
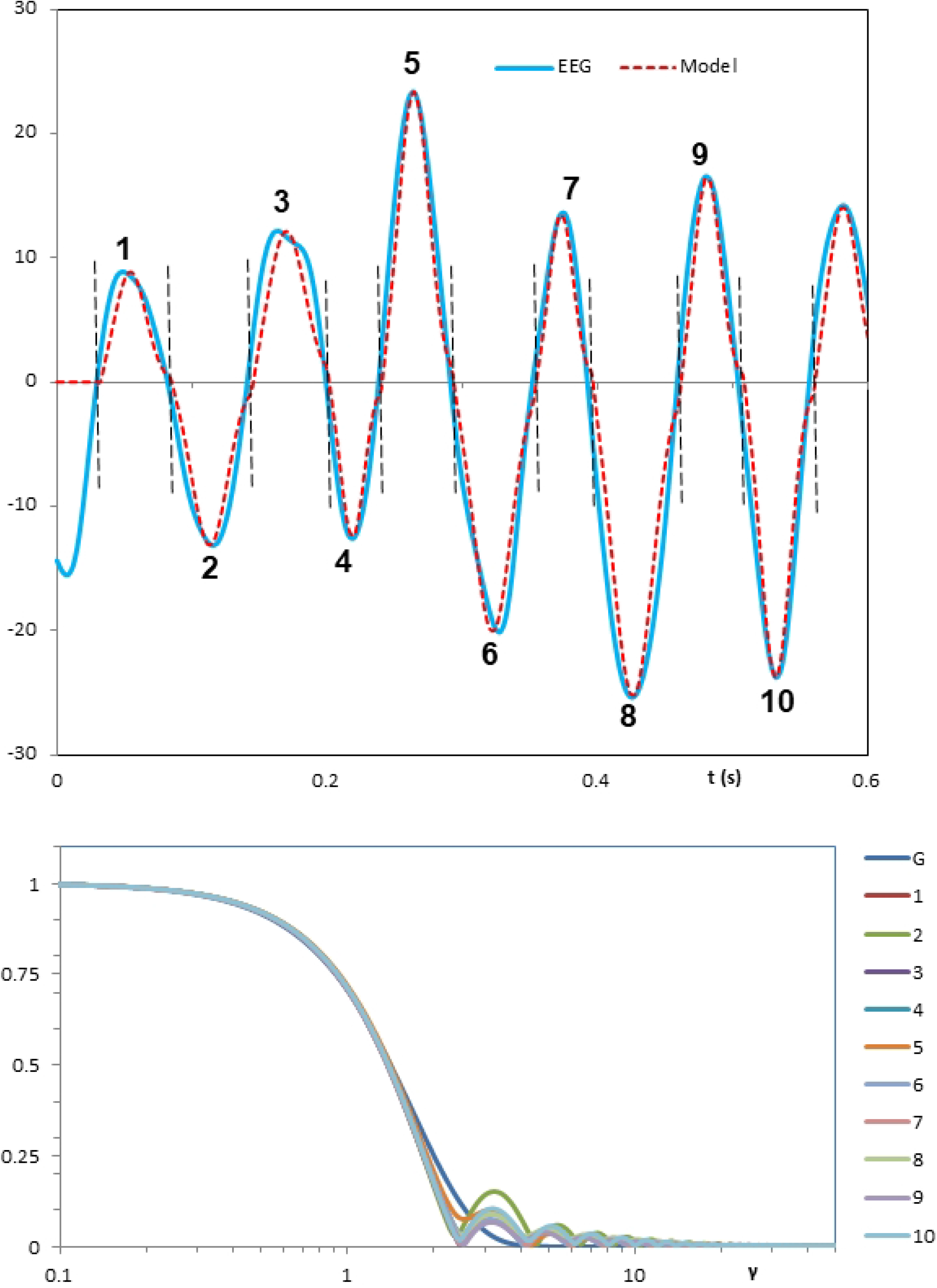
A typical example of the EEG time-frequency analysis. The y-axis units in the top panel are μV. The y-axis units in the bottom panel are μV·s.

The blue line in the top panel is the EEG recording divided into the ten segments by vertical dotted lines. Since the component overlap is minor, the zero-crossings have been taken as the segmentation points. Accordingly, a straightforward fragmentary decomposition was applied. Each piece of the signal between consecutive segmentation points has been transformed to the frequency domain using the SBF algorithm.

Computed amplitude spectra normalized by both the amplitude and frequency are depicted in the bottom panel. The blue line denoted by G is the Gaussian spectrum defined by Eq (11) (i.e. the model amplitude spectrum). Using the parameters σ and η defined for each segment in the frequency domain the model of the whole EEG signal in the form of the equation Eq(8) has been calculated. It is shown in the top panel by the red line.

An important aspect of similar procedures in the single trial ERP analysis is a need to take into account the component overlap. This problem is solved by using the HRFD. Typical results of the HRFD application for creation of an explicit model of a single trial ERP composed of multiple components is illustrated in Fig. 8.

**Fig 8.**
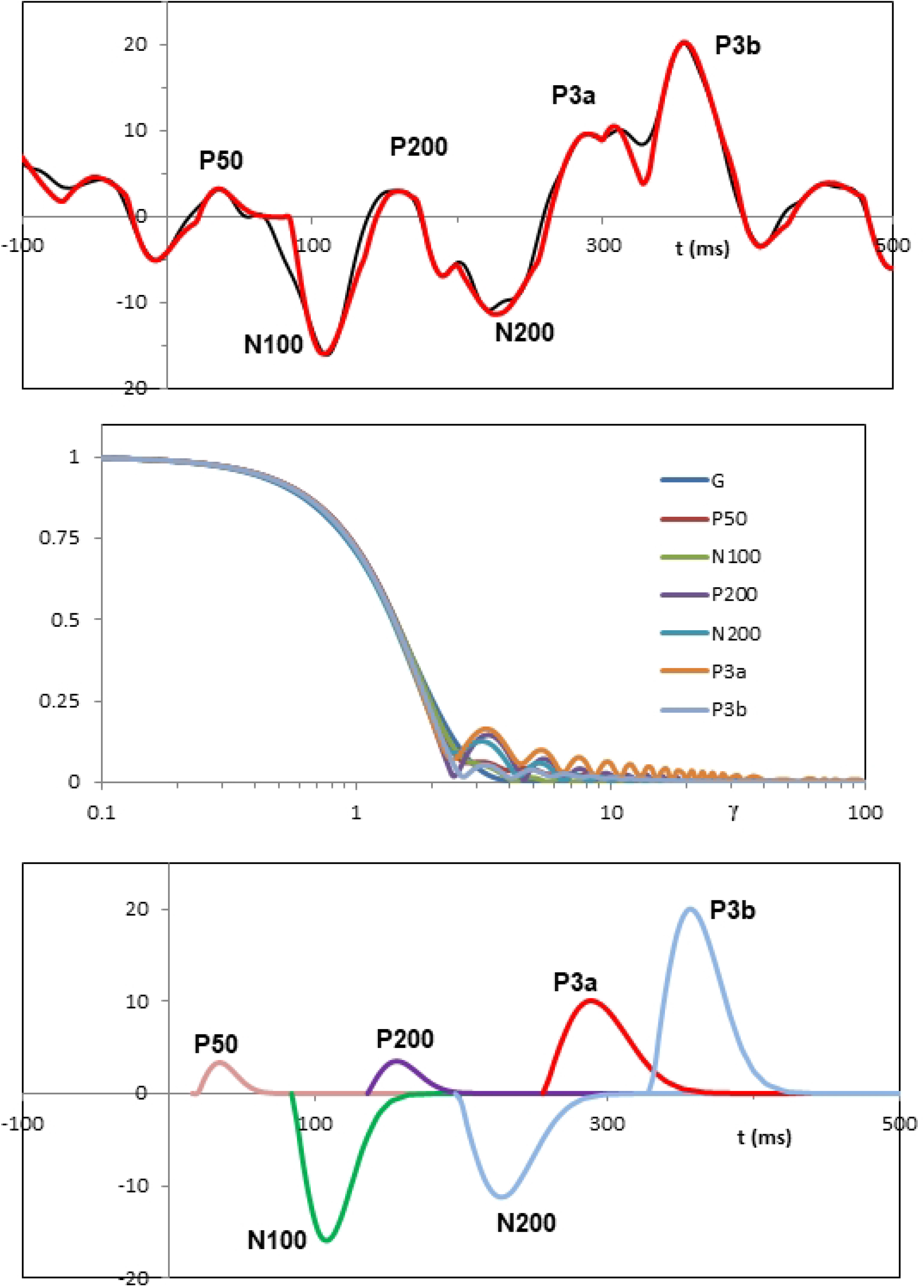
A typical model of single trial ERP from Cz recording site with its components identified by the HRFD.

The dynamics of the EEG signal shown in the upper panel on the interval from −100ms to 500ms is affected at t=0 by the application of the target auditory stimulus. This event is followed by a specific succession of positive and negative waveform deflections, the 6 species of which have been identified as conventional P50, N1, P2, N2, P3a and P3b late component ERPs.

Normalized amplitude spectra of these components are shown in the middle panel. The line “G” is the normalized amplitude spectrum defined by Eq(11). It appears as a limiting form of empirical spectra.

The sum of all identified HWFs is a model single trial ERP which is shown in the top panel by the red line. The models of separate ERP components are shown in the bottom panel.

A remarkably accurate match of the models to empirical curves is typical for the identification procedures of the HRFD. The key observation is a mutual coincidence of empirical amplitude spectra in a wide range of the relative frequency γ from 0.1 to 100.

An appealing feature of these results is that models of various EEG and ERP waveforms are obtained without requiring knowledge of the details of underlying cellular and molecular systems. This paradigm may be qualified as the phenomenon known in quantum theories as *universality* [32]. Conceptually, *universality* means that despite a profound diversity of complex dynamic systems observed in nature, particularly biology, their topology may have universal characteristics regarded as universal objects. Our methodology for the first time identifies such universal elements directly from the dynamics of EEG and ERP signals.

The universality indicates a stochastic nature of the mechanisms which produce the macroscale EEG waveforms and ERP components. The composition of the HWF as a sum of two shifted Gaussian functions suggests that the normal distribution governs transitions from the micro- to macro-scales.

The Kolmogorov-Smirnov tests were employed to examine this conclusion (i.e. a null hypothesis that the corresponding statistical regularities follow the normal distribution). The test has the advantage of making no assumption about the distribution of the samples (i.e. it is non-parametric and distribution free).

The ε (dimensionless extension ratio=F_B_/F_C_) has been selected as an adequate parameter for these tests because this single measure is sufficient to evaluate the fit of results.

Typical results illustrated in Fig 9 were obtained using single trial recordings from Fz, Cz and Pz cortical sites.

**Fig 9.**
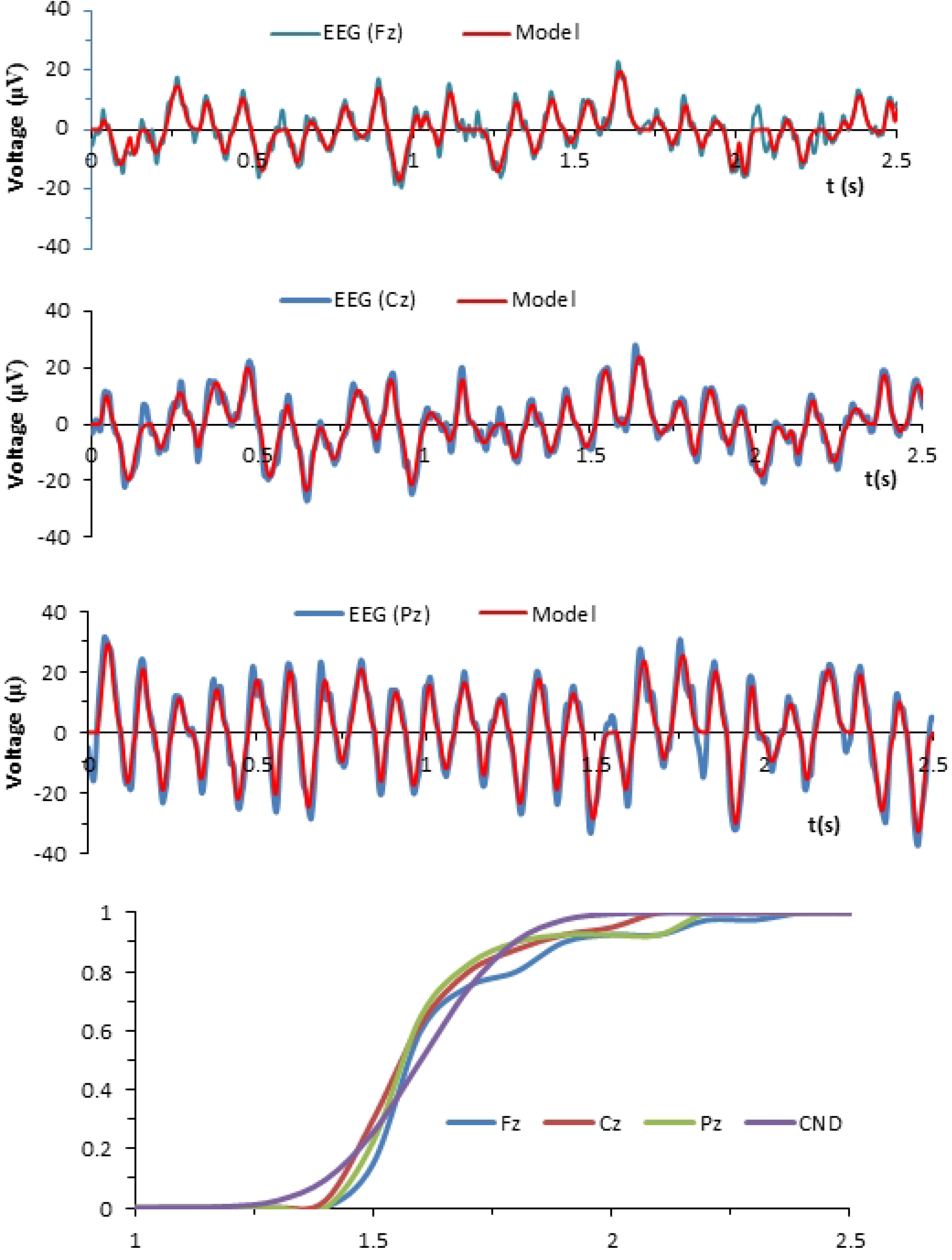
Upper panels show EEGs from Fz, Cz and Pz recording sites and remarkably accurate models calculated by the method of the HRFD. In the bottom panel the cumulative distribution functions of these processes are compared with the curve of cumulative normal distribution.

For each location the values of ε were collected from 20 different identified EHWs. The EHW was accepted as eligible for testing the D-statistics if the number of EHW’s samples was ≥8.

The means, standard deviations (SD) and D (K-S statistic) were as follows.

Fz: mean =1.756 (SD=0.205), D=0.106.
Cz: mean=1.712 (SD=0.152), D=0.076.
Pz: mean=1.721 (SD=0.182), D=0.079.

The blue lines in the upper panels show pertinent EEGs from Fz, Cz and Pz cortical sites. The red lines are the models calculated using the HRFD. The bottom panel shows the corresponding cumulative distribution functions normalized by sample size. The line denoted by CND is the cumulative normal distribution.

The greatest discrepancy between the CND and the empirical cumulative distribution, called the D-statistic, serves as a criterion to reject or accept the null hypothesis. Given that all calculated empirical cumulative distributions have been supported by equal numbers of ε (40 values of ε employed in the tests), the null hypothesis is rejected if D≥0.189 (5%).

The above estimates are well below this value and do not provide any reason to reject the null hypothesis. It is important to note the highly stereotypical character of the test results. The outcomes of multiple tests using the data from subjects from control and patient groups indicate the universality of introduced empirical distributions. Therefore, we can consider HWF an adequate universal model of ERP components.

### Conventional and synthetic grand mean ERPs

The analysis of specific features of ERPs in different groups of selected subjects conventionally uses grand mean averages. The left side panels in Fig 10 show grand mean average ERP waveforms calculated by conventional averaging for each group using 40 artefact free single trials from Fz, Cz and Pz middle sites. The histograms in the right-side panels show the elicitation rates of defined components. Taking the total number of single trials as 100%, the bars show the percentages of the sweeps in which components were identified, i.e. the frequencies with which different ERP components appear in single trials.

**Fig 10.**
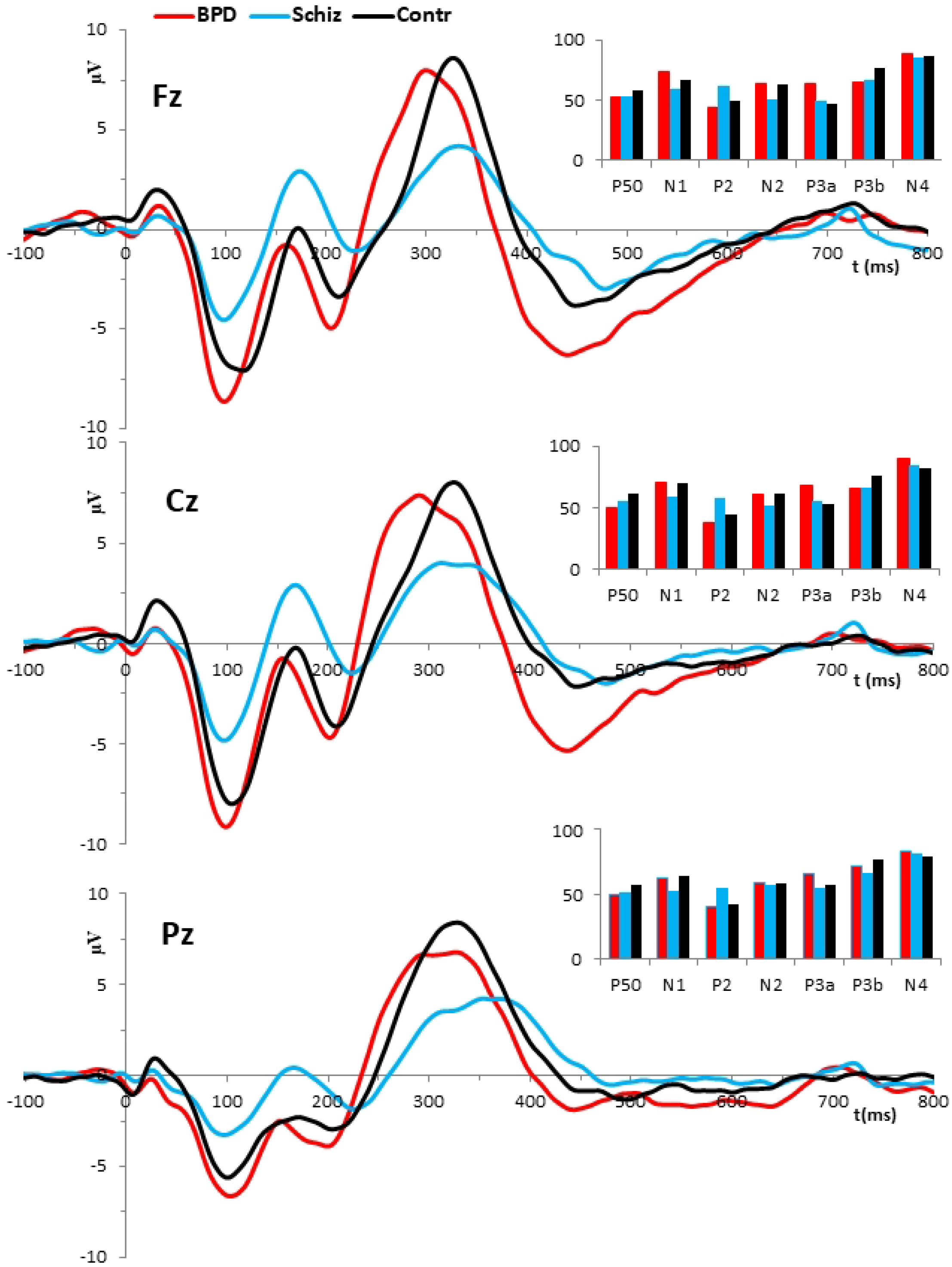
Grand mean averages obtained by conventional averaging.

One of the most investigated endogenous brain potentials in psychiatric research is the positive ERP wave at approximately 300ms peak latency range, denoted as P3. This component is clearly seen on all of these records as a monolithic positive waveform with notable broadening of the shape. In all recording sites, the P3s of schizophrenia patients show a decrease of peak amplitudes. For example, at Cz recording site the 8.03 μV amplitude of the P3 in control subjects is reduced in patients with schizophrenia to 4.02 μV. Such a trend is consistent with the general consensus that reduced P3 amplitude is one of the most replicable biological observations in schizophrenia [10]. Phenomenologically, it is generally accepted that P3 is created by the coordinated activity of multiple intracranial sources. With regard to the auditory oddball paradigm, there is a good deal of evidence that the P300 elicited by a target stimulus consists of two major components called the P3a and P3b [33]. These are independent and dissociable processes. The main distinguishing feature of the P3a is that it has a significantly shorter latency than P3b the peak latency of which is in the range of 300-400ms. The morphology of these potentials and their temporal overlap are easily recognizable in typical results of single trial analysis as exemplified in Fig 8.

However, the capacity of conventional averaging to identify the P3 as a composite of the P3a and P3b components is quite limited. It is particularly demonstrated by Fig 10 where the composite nature of the ERP in the 300ms range is obscured in conventional averages. For that reason, most previous techniques neglected the existence of the P3a and considered the ERP in the 300ms range as a single P3(00) component.

By contrast to conventional averaging, the HRFD identifies the whole complex of the late ERP components. Of crucial importance is that this methodology eliminates the temporal overlap of P3a and P3b components, allowing us to treat these ERP components as separate entities with different diagnostic features. The individuality and independence of the P3a and P3b components is supported by our finding that these components comprise 3 major types of activity pattern, both in the control and patient groups. This diversity of ERP patterns in the 300 ms latency range is exemplified in Fig 11.

**Fig 11.**
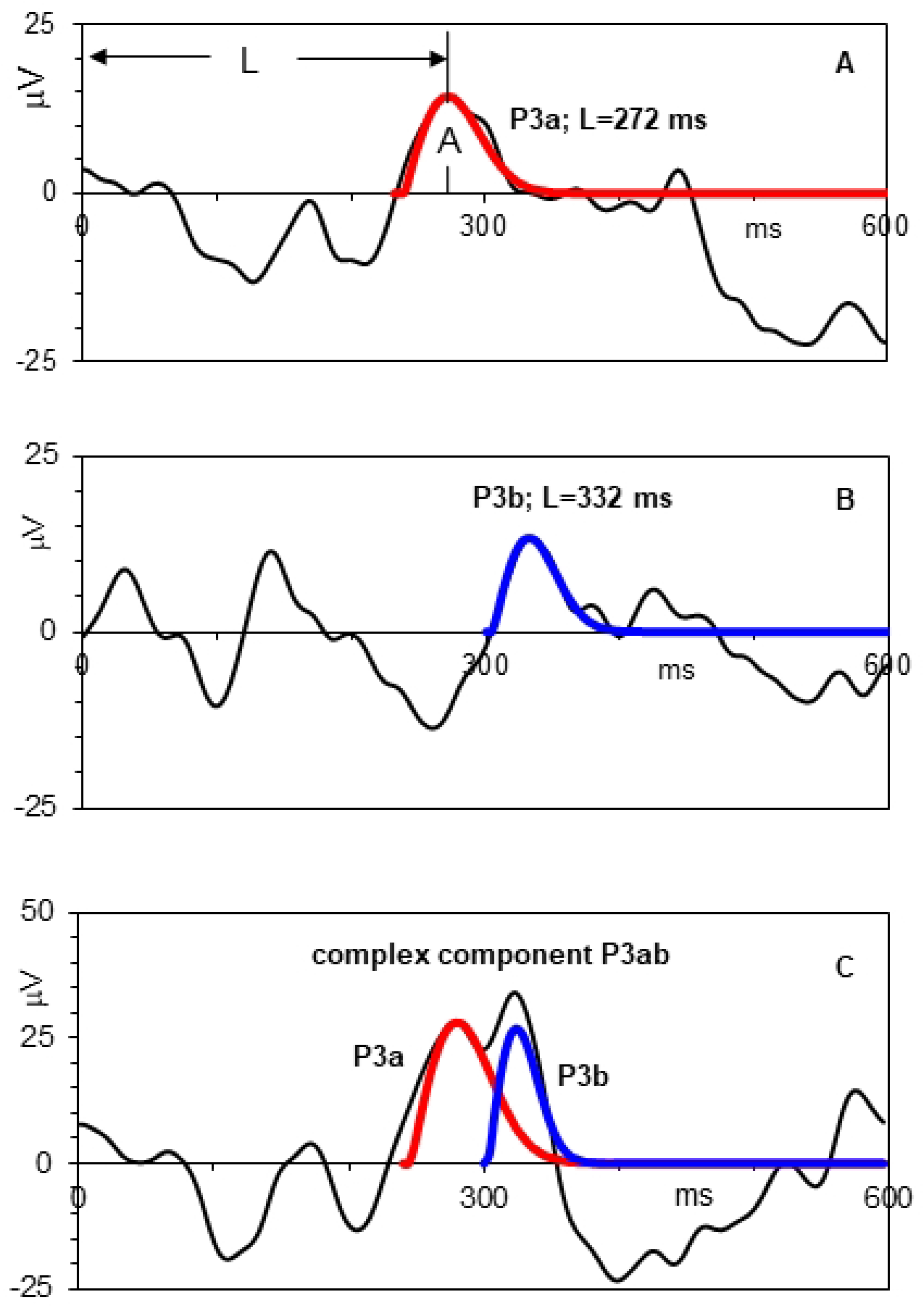
Various morphologies of the late positive component complex in single trial ERPs recorded in one and the same control subject from the Cz cortical site. The black lines show single trial ERPs induced by target stimuli applied at t=0. The colored lines are the models of the P3a and P3b.

The first two types in panels **A** and **B** are single trials with a sole type of subcomponent, i.e. a sole P3a and sole P3b, respectively. The third type in panel **C** is a complex component P3ab which develops as the superposition of P3a and P3b produced in one and the same single trial. These data show that there is no basis supporting the assumption of conventional averaging that the P3(00) is a monolithic component with an invariant pattern of activity. These aspects of the variability of ERP components, particularly the co-existence of P3a and P3b components, are obscured in conventional averages.

The HRFD considers a candidate event-related component as being not just a peak in the EEG waveform, but a whole deflection (positive or negative) with a particular shape defined by the set of 3 parameters. After the selection of “true” components using conditions 1-3, the procedure breaks down a single trial ERP into the sum of HWFs which provides a synthetic model of the single trial ERP. The sum of such models for selected components and cortical sites from selected groups of subjects provide synthetic grand mean averages.

Using the same original data as those supporting conventional grand mean averages in Fig 10, the synthetic grand mean ERPs are shown in Fig. 12.

**Fig 12.**
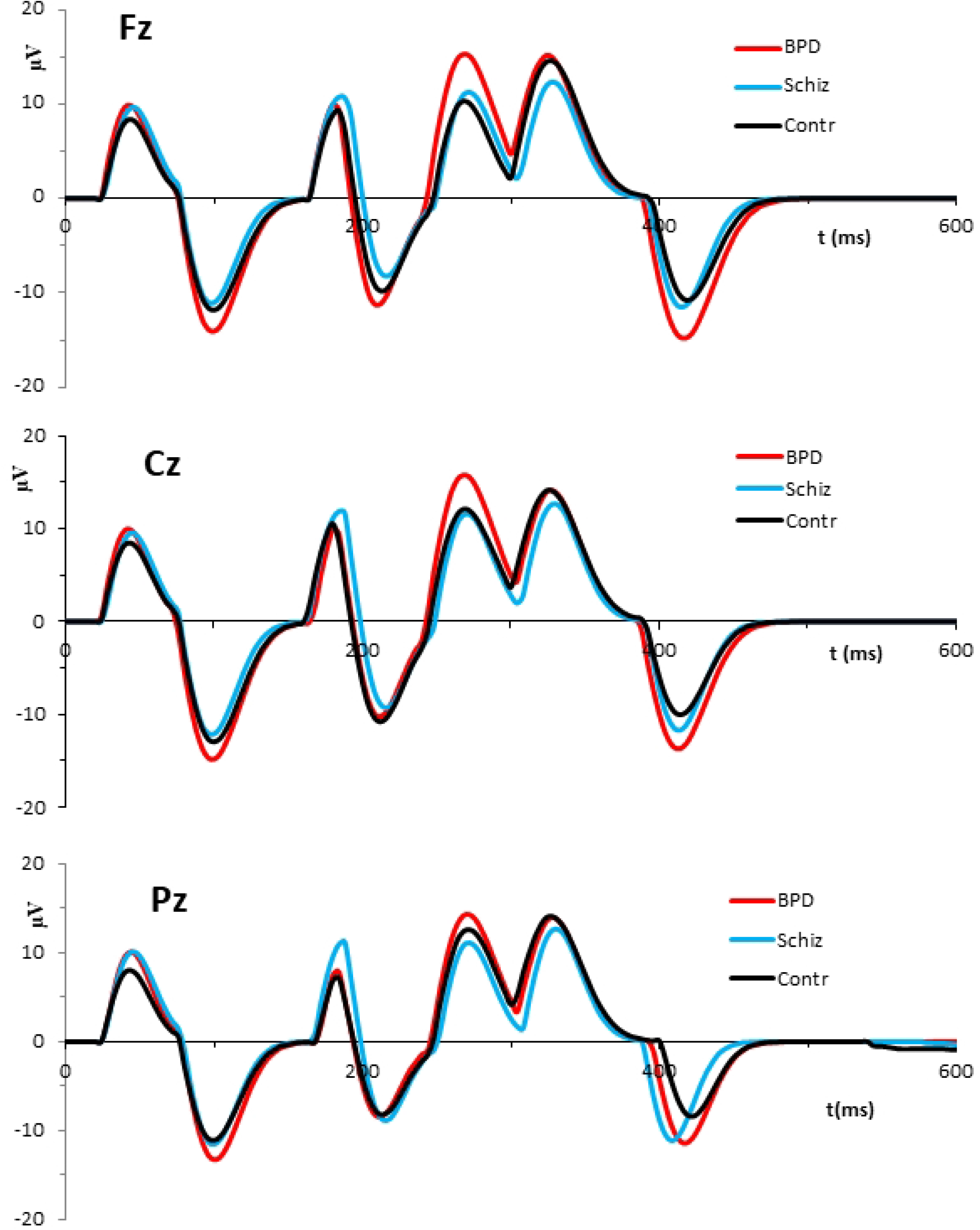
Synthetic grand mean averages for the same data as in Fig. 10.

We see marked differences between the results of the two methods of averaging. Compared with conventional averages, synthetic grand mean averages show significantly increased voltages. This indicates a crucial oversimplification in conventional ensemble averaging which assumes that ERP components of interest are present in all single trials. Consequently, the average is achieved by dividing the sum of all trials by the number of single trials. By contrast, SCA deals with actual numbers of identified components. For example, for the control group the numbers of identified components from the Cz recording site are: N_N1_=471, N_N2_=304, N_P2_=420, N_P3a_=353, and N_P3b_=505, where the subscript is the name of the component. Given P3b for example, this means that synthetic average is estimated by the SCA as the sum of 505 identified single trial P3b components divided by an exact number of identified components, i.e. the N_P3b_. For the same situation, conventional averaging divides the sum by 680 (i.e. the total number of single trials, 40 single trials for each of 17 subjects).

Another serious problem with conventional averaging is that significant temporal overlap of P3a with P3b creates numerous methodological complications for detection of these components in conventional grand mean averages. This situation is exemplified in Fig. 13 which compares the results of conventional (panel A) and selective component averaging (Panel B) using the data from the BPD and control groups.

**Fig 13.**
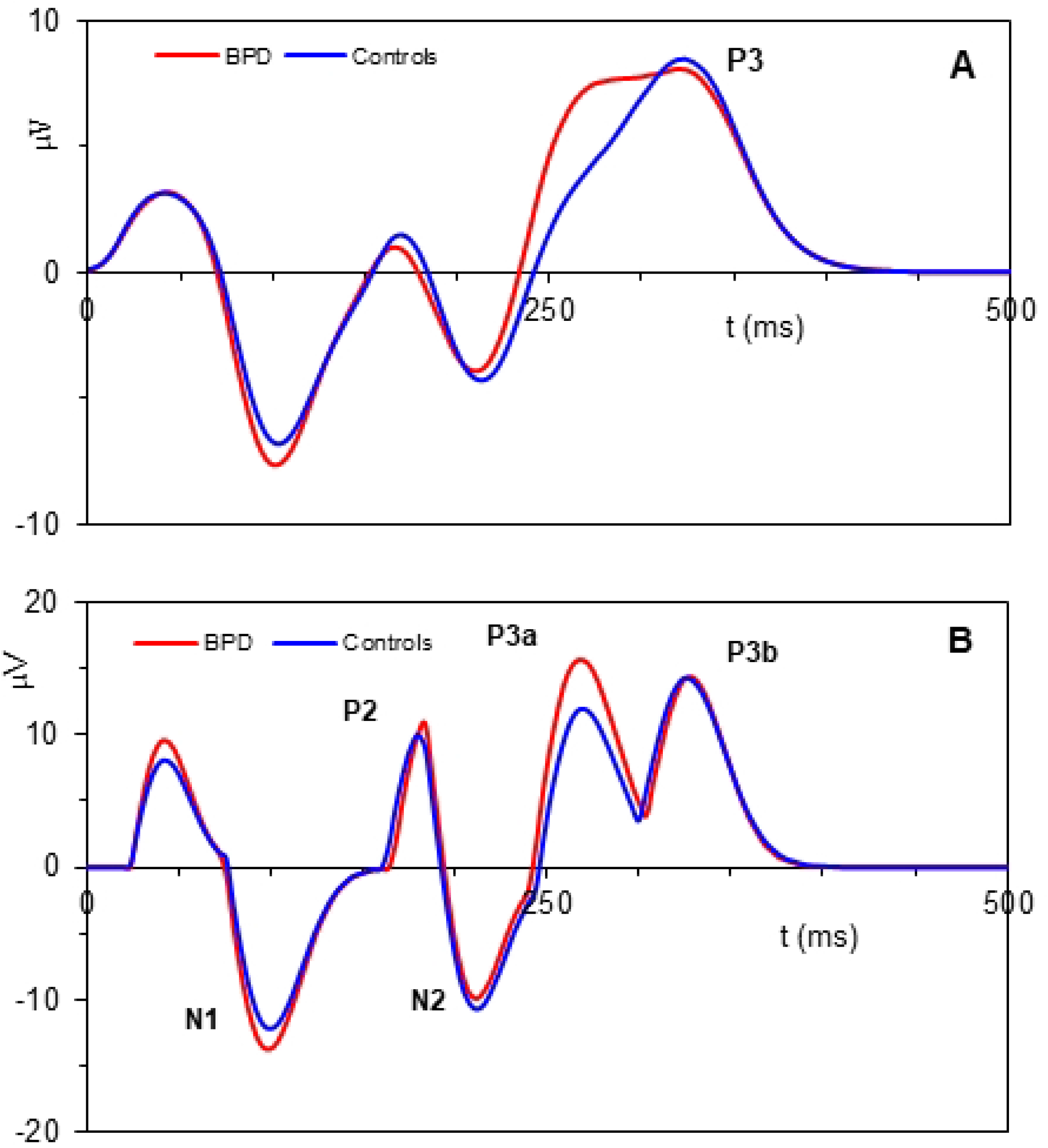
The panel A illustrates conventional grand mean averages in which P3 appears as a unitary component. By contrast, the synthetic grand mean average in the panel B reveals the P3a-P3b complex components in the 300 ms range.

The ambiguities involved in the interpretation of the ERP parameters from conventional grand mean averages are twofold. The first is that P3a and P3b components remain unknown. Secondly, the variable morphology of these components may affect the amplitude and latency parameters of the P3 from conventional averages in unpredictable ways. Without reliable methods for identifying the composite nature of late components of the ERP, many investigators evaluate complex average waveforms in the 300 ms latency range only for the maximum amplitude and latency, which assumes a monolithic average P3. These methodological difficulties and related simplifications may explain why the P3a has not been observed in ERP studies across different subject and patient populations as consistently as was the larger and more prominent P3b.

Reliable recognition of both the P3a and P3b components, unfulfilled by the previous methods, is achieved in our study through creation of a model-based approach to the component identification.

In the following sections we provide evidence that delineation of the P3a-P3b complex, instead of a single P3, has significant potential to enrich the diagnostic power of ERPs. Accordingly, the major focus is on features of P3a and P3b components. The grand mean averages of these components are shown in Fig. 14.

**Fig 14.**
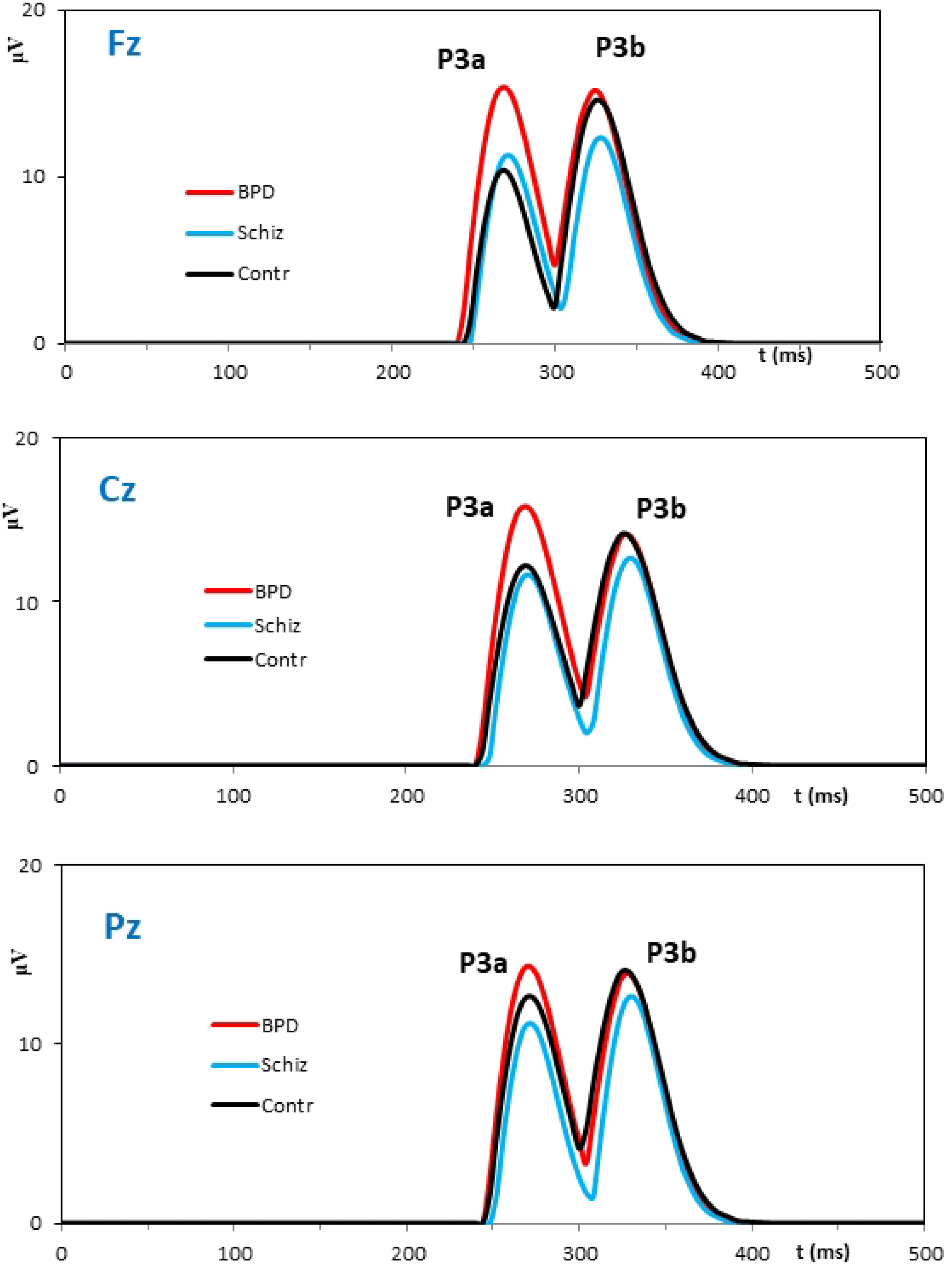
Grand mean averages of the P3a and P3b components.

The major parameter which defines single trial EEG deflection as a P3a or P3b component is the peak latency. The values of this parameter for middle sites in the 3 groups are listed in the table 1.

**Table 1.**
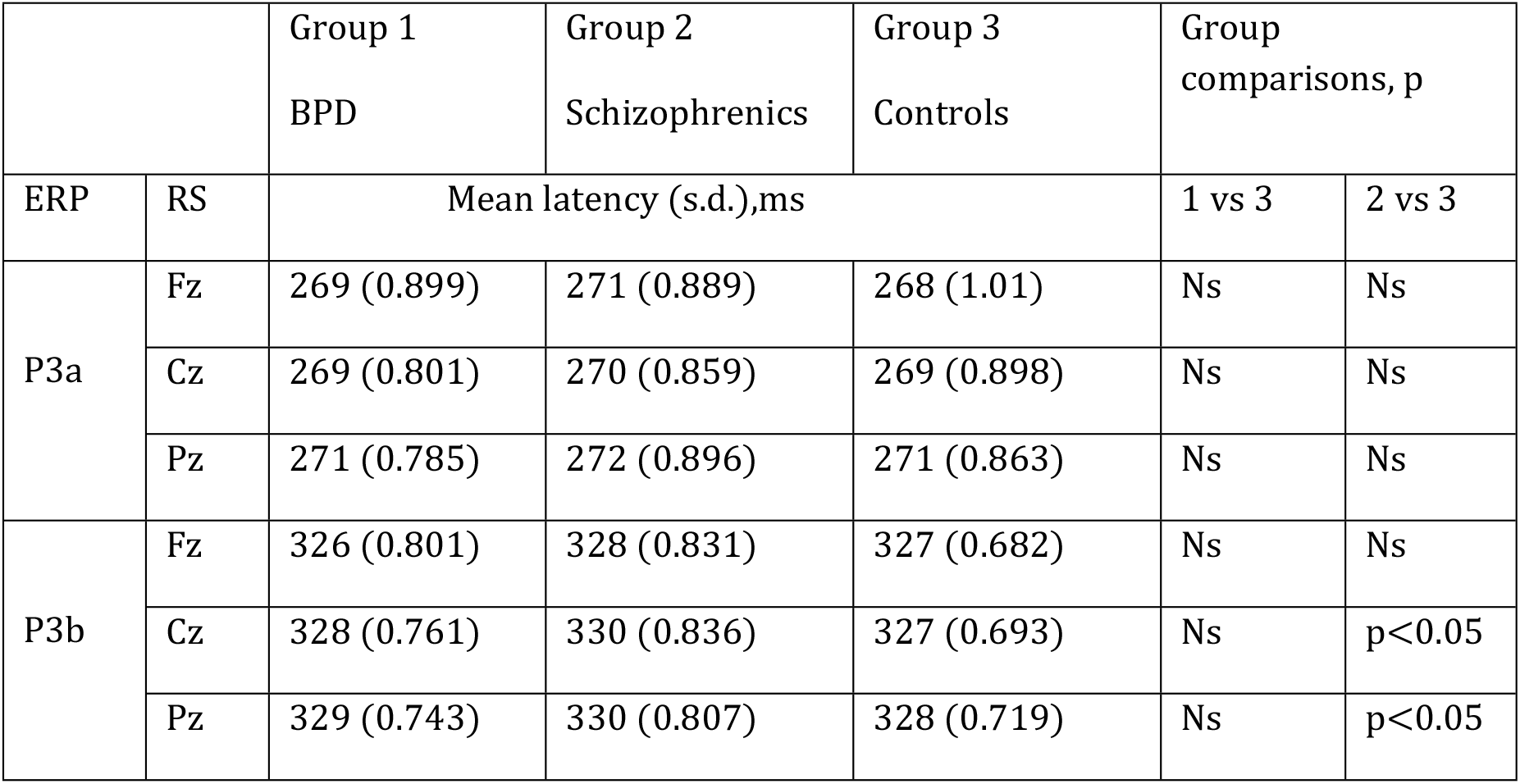
Shows mean latencies and standard deviations (s.d.) in midline electrode sites of subjects from the two patient groups (BPD and schizophrenics) and the group of controls.

|  |  | Group 1<br>BPD | Group 2<br>Schizophrenics | Group 3<br>Controls | Group comparisons, p |  |
| --- | --- | --- | --- | --- | --- | --- |
| ERP | RS | Mean latency (s.d.),ms |  |  | 1 vs 3 | 2 vs 3 |
| P3a | Fz | 269 (0.899) | 271 (0.889) | 268 (1.01) | Ns | Ns |
|  | Cz | 269 (0.801) | 270 (0.859) | 269 (0.898) | Ns | Ns |
|  | Pz | 271 (0.785) | 272 (0.896) | 271 (0.863) | Ns | Ns |
| P3b | Fz | 326 (0.801) | 328 (0.831) | 327 (0.682) | Ns | Ns |
|  | Cz | 328 (0.761) | 330 (0.836) | 327 (0.693) | Ns | p<0.05 |
|  | Pz | 329 (0.743) | 330 (0.807) | 328 (0.719) | Ns | p<0.05 |

The justification of the closeness in values of peak latencies may be attributed to the fact that our methodology of the HRFD eliminates the component overlap which introduces uncontrolled errors in the estimated parameters of average ERPs. The number of trials selected for HRFD can be found in the tables 2 and 3. A visual impression of the morphological similarities is demonstrated by the graphs in Fig. 14.

**Table 2.**
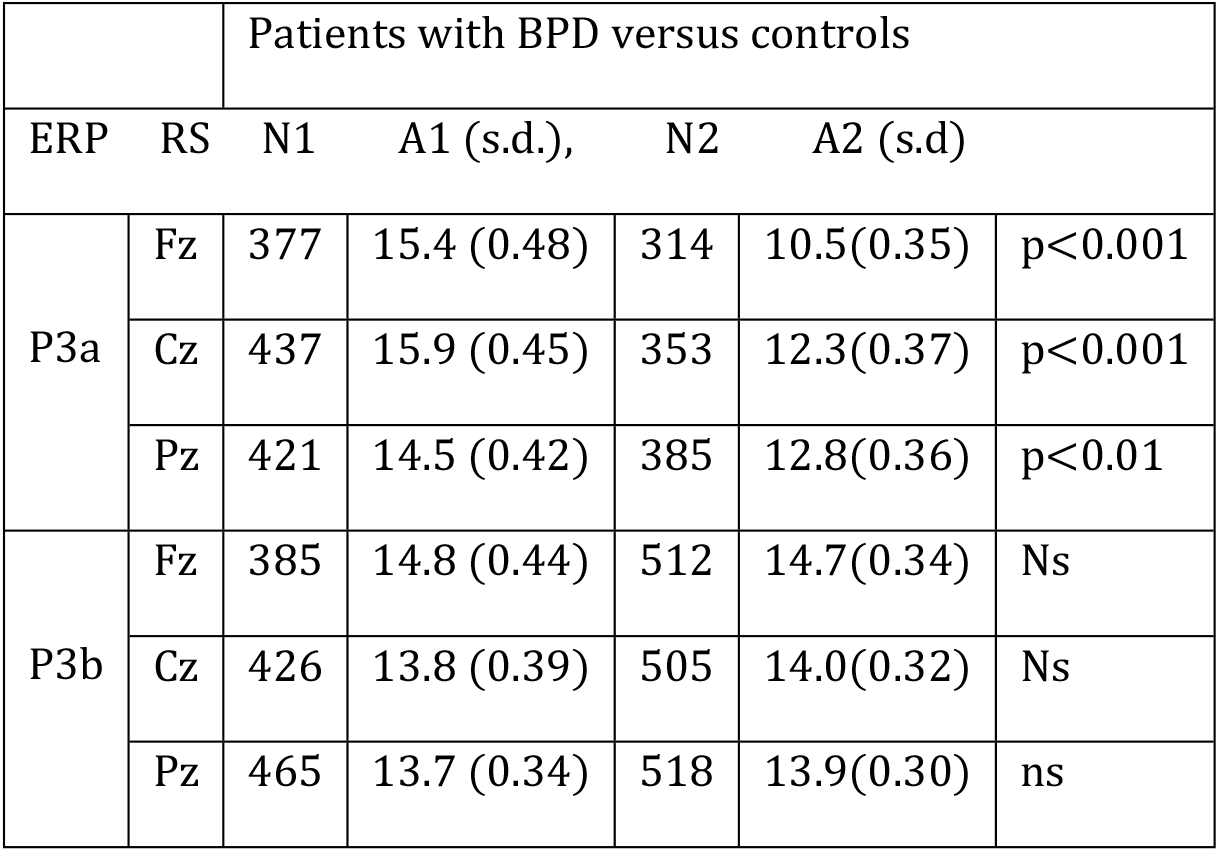
Compares the peak amplitudes of the P3a and P3b components for Fz, Cz and Pz recording sites in the subjects from the BPD and control groups. RS denotes recording site and N is the number of single trials selected for HRFD. The A1 and A2 are the peak amplitudes from the BPD and control groups, respectively.

**Table 3.**
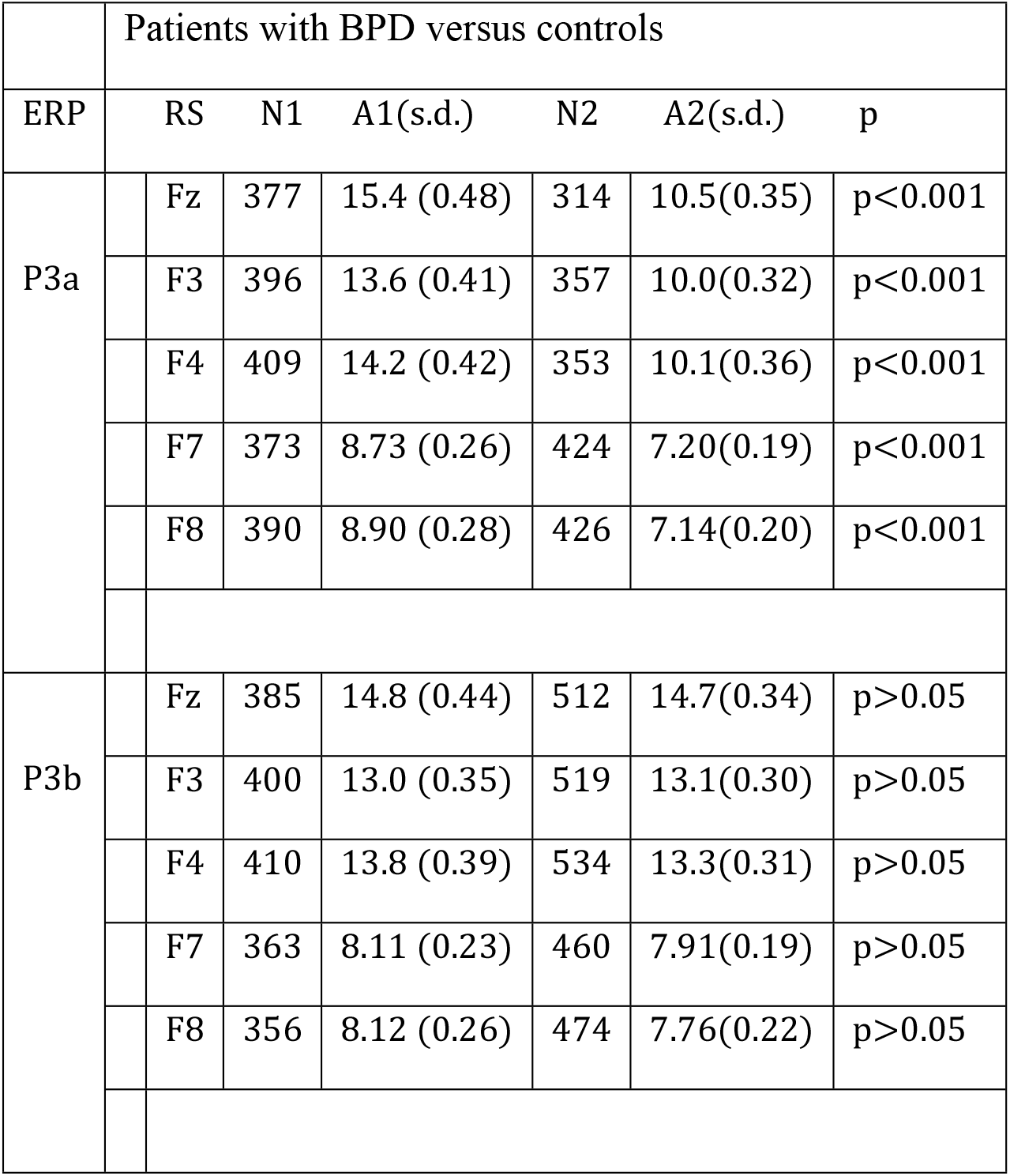
Comparison of peak amplitudes of P3a and P3b at frontal electrode sites. These sites are illustrated in Fig 15.

| Patients with BPD versus controls |  |  |  |  |  |  |
| --- | --- | --- | --- | --- | --- | --- |
| ERP | RS | N1 | A1(s.d.) | N2 | A2(s.d.) | p |
| P3a | Fz | 377 | 15.4 (0.48) | 314 | 10.5(0.35) | p<0.001 |
|  | F3 | 396 | 13.6 (0.41) | 357 | 10.0(0.32) | p<0.001 |
|  | F4 | 409 | 14.2 (0.42) | 353 | 10.1(0.36) | p<0.001 |
|  | F7 | 373 | 8.73 (0.26) | 424 | 7.20(0.19) | p<0.001 |
|  | F8 | 390 | 8.90 (0.28) | 426 | 7.14(0.20) | p<0.001 |
| P3b | Fz | 385 | 14.8 (0.44) | 512 | 14.7(0.34) | p>0.05 |
|  | F3 | 400 | 13.0 (0.35) | 519 | 13.1(0.30) | p>0.05 |
|  | F4 | 410 | 13.8 (0.39) | 534 | 13.3(0.31) | p>0.05 |
|  | F7 | 363 | 8.11 (0.23) | 460 | 7.91(0.19) | p>0.05 |
|  | F8 | 356 | 8.12 (0.26) | 474 | 7.76(0.22) | p>0.05 |

### P3a and P3b in BPD patients compared with controls

The data from the table 1 show that the differences in the peak latencies of P3a and P3b from the group of BPD patients compared with controls are non-significant for all middle recording sites. In contrast, the data from the table 2 show statistically highly significant increases of P3a peak amplitudes at Fz, Cz and Pz middle sites in BPD patients.

An important feature of these estimates, also evident in Fig 14, is a significant increase of P3a amplitudes in the group of BPD patients. In contrast, estimates of the P3b peak amplitudes didn’t reveal significant inter-group changes.

The differences between the values of the P3a peak amplitudes in patient and control groups show increases from Pz to Fz sites. These are: Pz – 1.7 (μV), Cz −3.7 (μV), Fz −5.0 (μV).

This trend indicates that the frontal areas of the brain are altered in BPD. Additional support for this tendency is provided by the data in Table 3 with similar comparisons of the peak amplitudes from Fz, F3, F4, F7 and F8 frontal electrode sites.

### P3a and P3b in patients with schizophrenia

The results of inter-group measurements of the P3a and P3b peak amplitudes in patients with schizophrenia versus controls are presented in Table 4.

**Table 4.** Peak amplitudes of P3a and P3b components for Fz, Cz and Pz recording sites in the subjects from the schizophrenics and control groups.

| Patients with schizophrenia versus controls |  |  |  |  |  |  |
| --- | --- | --- | --- | --- | --- | --- |
| ERP | RS | N1 | A1 (s.d.) | N2 | A2 (s.d.) |  |
| P3a | Fz | 315 | 11.4 (0.37) | 314 | 10.5(0.35) | Ns |
|  | Cz | 373 | 11.8 (0.35) | 353 | 12.3(0.37) | Ns |
|  | Pz | 370 | 11.3 (0.34) | 385 | 12.8(0.36) | Ns |
| P3b | Fz | 430 | 12.3 (0.34) | 512 | 14.7(0.34) | p<0.001 |
|  | Cz | 441 | 12.7 (0.33) | 505 | 14.0(0.32) | p<0.01 |
|  | Pz | 442 | 12.7 (0.35) | 518 | 13.9(0.30) | p<0.01 |

RS denotes the recording site. N1 and N2 are the numbers of single trials selected from the group of patients with schizophrenia and controls, respectively. The A1 and A2 are peak amplitudes from schizophrenia and control groups respectively.

The major outcomes are twofold. First, the P3a amplitudes in schizophrenia patients do not show significant changes compared with the data from the group of control subjects. Second, the P3b amplitudes in schizophrenia patients are significantly reduced at all middle electrode sites.

The grand mean averages in Fig. 14 give a clear picture of these patterns.

## Discussion

The main findings reported in this paper depend critically on the methodological innovation of EEG and ERP analysis using the probabilistic methods of quantum theories. This novel approach provides a means of bridging the macro-scale EEG and ERP signals with the underlying molecular events of ion transport at the micro-scale.

We first discuss the micro-scale results, the main aspect of which is the relocation of elementary bioelectric sources of the EEG signal from the cellular to the molecular level. Instead of the continuous time membrane potentials implemented in previous models, the elementary cortical sources of electricity are now allocated to ions, positively and negatively charged particles, the size and stochasticity of which conform in their attributes to quantum mechanics. The vanishingly small role of individual charges in the generation of macroscopic scale EEG signals reduces the problem to the study of the behavior of large numbers of random variables underlying the phenomenon in question. This is realized through the predictions of the central limit theorem.

The idea of EEG interpretation using probabilistic notions was previously put forward by Elul [34]. Based on analysis of the synchronization of EEG sources, Elul proposed that the evolution of brain waves may be governed by statistical regularities following from the central limit theorem. Thus, the EEG waveform may simply be accounted for as a normally distributed output resulting from the combination of the activity of many independent neuronal generators.

This hypothesis has never been supported by an adequate empirically-testable mathematical theory. However, it raised the possibility that “gaussianity” may be the most promising explanatory feature of ERPs. The straight realization of this approach is employment of the normal distribution as a model of N1, P2, N2 and P3 components of average ERPs [35]. One of the major factors that introduce significant inconsistency between the Gaussian function and real ERP waveforms is that normal distribution is a bell-shaped symmetrical function defined on an infinite time scale while the ERP component is a transient process the start of which is locked to the cognitive event with different rates of rise and decay. To take into account a steeply rising left flank and a slowly decreasing right flank the convolution of an exponential and a normal distribution (ex-Gauss function) has been used for quantifying the P3 component in average ERPs [36]. The gamma function is quite flexible and might be one of a variety of possible shapes that fit the component waveforms of average ERPs. However, the ex-Gauss function differs from predictions of the central limit theorem and cannot serve as a universal model of ERP components. In this context, the fundamental point of our approach is consideration of each ERP component as a statistical limit of the underlying microscale processes, the appearance of which, on the global scale, is governed by the central limit theorem. The prediction of this theorem is the normal distribution (i.e. the Gaussian function).

A single Gaussian function defined on an infinite scale is not a proper approximation of the ERP component, which is a transient starting from the moment of activation of the underlying cellular machinery by a cognitive event. This feature of ERP components is adequately described in our theory by Eq (2) composed of a sum of two shifted Gaussian functions the profile of which appears as an adequate form of the shape of the ERP component. We regard this paradigm as an indication of two ensembles of elementary charges which we call primary and secondary particle populations.

Physically, cell membranes, which separate intracellular from extracellular space, play a crucial role in the creation of the microscale model. Due to their high electrical resistance, membranes act as a border which prevents intracellular ions from noticeably changing extracellular field potentials. This means that extracellular ions appear to be the source of the global scale EEG and ERP. The impact of a single ion to the field potential is vanishingly small. Therefore, the changes of macroscale potentials are considered as cumulative effects produced by the transport of ions during synchronized activation of ensembles of closely located excitable cells.

In keeping with this, the modelling tools are changed from the deterministic equations of classical physics to the probabilistic formalism of non-homogenous BDPs with time dependent rates of birth and death. We define the specific aspects of this amalgamation of deterministic and stochastic factors on the microscopic scale as the transient deterministic chaos.

We now turn from microscale events to the macroscale phenomena which they produce in the form of EEG and ERP waveforms. The major achievement provided by the methodological innovations of single trial ERP analysis and the creation of synthetic grand mean averages is reliable identification of the P3a and P3b endogenous potentials in the 250-450 latency range. In general, on the basis of conventional averaging, the ERPs in this region have been conceptualized as the P300 arising from a single neural generator. This concept has shaped virtually every aspect of P300 research, including the way it is used in clinical studies. In this context, P300 has been used as an aid to diagnose neuropsychiatric disorders, sub-types of disorder and to evaluate the effects of medication on aspects of cognition. In reference to higher-order cognitive processes, no other endogenous potential has received as much attention from researchers in the last two decades as the P300. However, converging evidence from a number of experimental and clinical studies has made it evident that significantly different combinations of neural generators contribute to the P300 activity elicited by different combinations of experimental variables. This means that the P300 is not a monolithic component.

The most commonly recognized subcomponent in “oddball” tasks is the classical P3b which has a parietal maximum scalp distribution and a peak latency of 300-400ms. This is often preceded by a subcomponent, identified as a new component [33]. This component was labeled “P3a”, to distinguish it from the classical Suttonian P300 [37] which was re-labeled “P3b”. These P3 subcomponents usually overlap in time, making it difficult to recognize them in the time course of average ERPs. A specific problem is that ERP waveforms are not measured in single trials and then averaged, but are measured only once, in the average curve. This leads to the loss of crucial information about the morphologies of P3a and P3b components and the rates with which they respond to cognitive stimuli. Hence, the potential for the ERP to provide objective electrophysiological measures of cognitive variables to a large degree depends on our ability to analyze ERP component composition directly from single trial records. Though various filters and templates have been employed, comprehensive single trial ERP analysis has not been achieved using existing methods.

As far as we are aware, our study is the first to provide an empirically testable, adequate model of single trial ERP components. This methodological innovation allows us to eliminate the temporal overlap of P3a and P3b components in single trials and significantly improve the accuracy of the corresponding amplitude and latency parameters. An important and somewhat unexpected finding is the stability of the latencies of both the P3a and P3b components. As the data from the Table 1 indicate, in the control and BPD patient groups the mean peak latency of the P3a at all midline sites is in the range from 268 to 271ms. For P3b the range is from 326 to 329. For both the P3a and P3b no statistically significant inter-group differences were found.

In schizophrenia patients the inter-group differences in the peak latencies of P3a (the range from 268 to 272 ms) are non-significant. For P3b (the range from 327 to 330 ms), statistically significant differences in comparison with the control group were documented at Cz and Pz sites.

In contrast to the latency findings, we found profound changes in the peak amplitudes of the P3a and P3b in both patient groups. A critical finding is that there is a qualitatively different character of ERP morphology in the groups of patients with schizophrenia and BPD, suggesting functional differences in the underlying neuropathological processes.

The main distinguishing feature of ERP changes in BPD patients is an abnormal increase of the P3a peak amplitudes compared with control subjects. Table 2 shows a statistically highly significant increase of P3a peak amplitudes at Fz, Cz and Pz middle sites in BPD patients.

The data from Table 3 and Fig 15 demonstrate the frontal origins of this abnormality. Failure of inhibitory control may be the factor that accounts for the increase in amplitude of the P3a. In a wider neuro-psychiatric context, our recent aetiological model of BPD suggests that impairment of inhibitory control in prefrontal networks may underlie the disorder [38]. This P3a enlargement gives support to Meares’ hypothesis that the sense of disconnectedness, or disintegration, that is a core phenomenon of BPD, is the outcome of deficient higher order inhibitory function. This hypothesis is based upon a Jacksonian model of dissociation, developed further in a current texts on the subject [39, 40].

**Fig 15.**
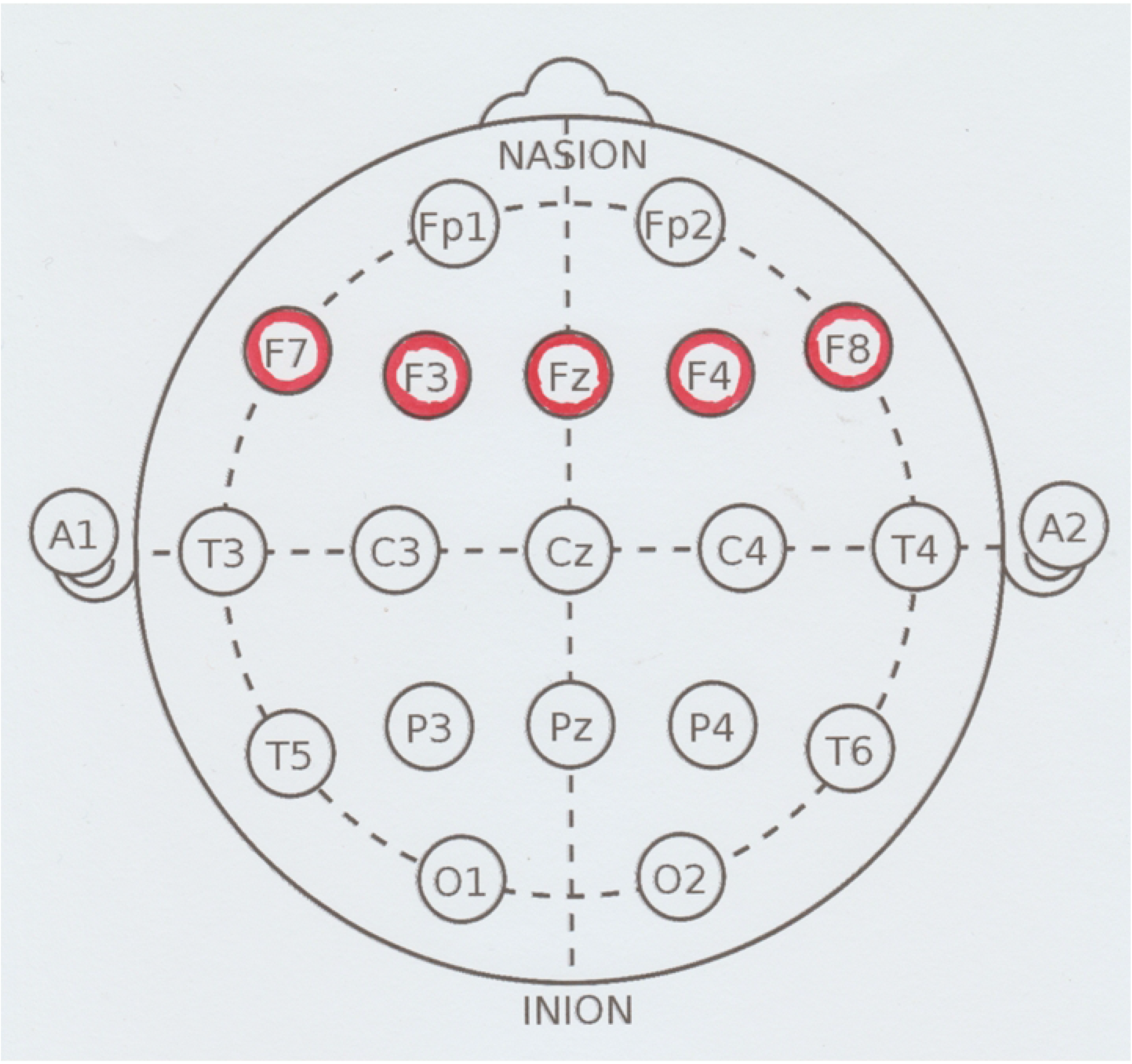
Topographic map of electrode sites. The circles rendered in red denote the sites of cortical electrodes at which the peak amplitudes of the P3a from the group of BPD patients have shown a statistically highly significant increase of peak amplitudes.

Our data demonstrate, as the main distinguishing feature of ERP changes in patients with schizophrenia, an abnormal decrease of the peak amplitudes of P3b while the voltage of P3a is not affected by this pathology. The reduction in amplitude of the average P3 from the standard auditory oddball paradigm is one of the most replicable biological observations of schizophrenia that is present regardless of medication status [10].

Our study is consistent with the previous single-trial analyses of P3 in patients with schizophrenia, in that the voltage reduction of the average P3 depends on two factors [10]. The first factor is a replication of previous analyses, where the voltage reduction of P3 components is identified in single trials. This indicates a reduced impact of microscale sources on the macroscale P3. A second factor, newly identified in this study, is the demonstration of a decreased number of single trials which contain the P3b component. The techniques of single trial ERP analysis employed in previous studies were limited in terms of capturing the P3a and P3b sub-components of the P3. The investigation of spatiotemporal distributions of single trial P3a and P3b using the methodological innovation of HRFD suggests a distinct character of these components. The major properties of the average P3 resemble single trial P3b. However, the temporal overlap of the P3a and P3b components and their changing patterns from trial-to-trial cause unpredictable changes of the latency and amplitude parameters of average P3s.

The P3a has been studied in patients with schizophrenia much less frequently than P3b. Usually, lower or unchanged amplitudes of P3a have been described in schizophrenia [41]. Consistent with these findings, our study did not reveal significant changes in the voltages of P3a.

A main inference from these most important results is that BPD and schizophrenia are physiologically different. Since they are disorders which have in common the identifying feature of disconnectedness, it is sometimes argued that they are the same. However, this study demonstrates an abnormality of P3a which is clearly defined in BPD but absent in schizophrenia.

